# miR-133a and miR-338-3p Shape Neural Crest Derivatives in Zebrafish

**DOI:** 10.1101/2025.06.06.658297

**Authors:** Steeman Tomás José, Weiner Andrea Maria Julia, Rubiolo Juan Andrés, E Sánchez Laura, Calcaterra Nora Beatriz

## Abstract

Neural crest (NC) cells are a transient, multipotent cell population that contributes to the formation of diverse tissues during vertebrate development. While numerous transcription factors and signalling pathways regulate NC specification and differentiation, the role of microRNAs (miRNAs) in these processes remains underexplored.

In this study, we employed a double-transgenic zebrafish model (Tg(*sox10*:eGFP, *sox10*:mRFP)), in combination with fluorescence-activated cell sorting (FACS) and RNA sequencing, to identify miRNAs enriched in NC cells and assess their role in the development of NC derivatives. Given the parallels between NC development and tumorigenesis, we focused on a subset of miRNAs previously implicated in cancer progression.

Functional assays revealed that overexpression of miR-133a and miR-338-3p resulted in significant craniofacial cartilage malformations and a reduction in melanophore numbers, correlating with the downregulation of key developmental genes, such as *sox9b*, *sox10*, and *runx3*. Reporter assays further demonstrated that miR-133a and miR-338-3p directly target the 3’ untranslated region (3’UTR) of *sox9b*, supporting their role in translational repression and mRNA degradation. Additionally, miR-338-3p overexpression increased the overall NC cell population, suggesting a regulatory function in cell proliferation.

These findings offer new insights into the molecular mechanisms governing NC development and highlight functional parallels between miRNA-mediated NC regulation and tumour biology. Given that miR-133a and miR-338-3p are known tumour suppressors, their roles in NC differentiation and proliferation may reflect conserved pathways underlying both embryogenesis and oncogenesis. This study enhances our understanding of miRNA function in vertebrate embryogenesis and underscores their potential relevance in neurocristopathies and NC-derived cancers.

## 1 Introduction

During vertebrate embryogenesis, the development of organs depends on the coordinated formation, remodelling, and, in some cases, the regression of various tissues and structures. Among these, the neural crest (NC) is a transient, multipotent cell population that emerges early in development and plays a pivotal role in generating diverse cell lineages. Initially located dorsally to the neural tube, NC cells (NCC) undergo extensive proliferation followed by epithelial-to-mesenchymal transition (EMT) which facilitates their delamination and migration [1,2]. Subsequently, these cells differentiate into a variety of derivatives, including neurons, glial and Schwann cells, craniofacial chondrocytes, pigment cells, and endocrine cells [3]. Defects in NC development are linked to neurocristopathies and certain cancers. Thus, elucidating the molecular mechanisms governing NC formation, migration, and differentiation is critical for understanding the aetiology of these disorders and for advancing therapeutic approaches [4,5]. NC development is regulated by a complex network of transcription factors (TFs) and signalling molecules, organized in a gene regulatory network (GRN) [6] that operates in a spatiotemporal manner. In zebrafish, NC formation is initially guided by extrinsic factors, such as the BMP and WNT pathways [7,8], which activate NC-specific genes such as *foxd3*, *snai1/2,* and *sox10* [9]. During EMT, the expression of new TFs such as *twist1a/b* facilitates the switch from epithelial to mesenchymal cadherins, among other changes, enabling migration [6,10,11]. Once NCC reach their destination, additional external signals induce further regulatory changes specific to the GRN of each derivative.

NCC give rise to pigment cells, such as melanophores, iridophores and xanthophores in zebrafish. Melanophores are responsible for black pigmentation and emerge at 24 hours post-fertilization (hpf); their appearance depends on the expression of *sox10* and *pax3/7*, which activate the master gene *mi8a [*. Mitfa, in turn, induces the expression of melanogenic genes such as *dct, tyr*, and *pmel* [13,14]. Epigenetic regulators like HDAC1 appear to repress alternative NC fates [14], favouring melanophores differentiation. Iridophores, responsible for iridescence through guanine crystals, appear at around 72 hpf [12,15] and follow a similar regulatory pathway but diverges through Tfec-mediated activation of TFs *ltk* and *sox10* [16–18], among other factors. NCC also contribute to craniofacial chondrocytes development [19,20] and skull formation in zebrafish. In this process, *sox9* paralogs (*sox9a* and *sox9b*) [21,22], modulated by *runx3* and *egr3*, regulate chondrocyte stacking and proliferation [23]. Sox9a/b, in conjunction with Sox5/6, activate the expression of *col2a1a,* a key collagen-coding gene in cartilage, and *agc1,* a marker of cartilage differentiation. Given the complexity of craniofacial development, disruptions in these regulatory pathways can contribute to congenital disorders.

Recent studies have highlighted the critical role of microRNAs (miRNAs) in NC development [24,25]. These 22nt endogenous, non-coding RNAs regulate mRNA stability and transcription [26] through the miRISC complex, which includes the RNAse Argonaut. The miRISC complex targets mRNA by binding to a specific “seed region” of the miRNAs [27,28], typically within the mRNA’s 3’UTR, and each miRNA family has an average of 300 binding sites under selective pressure [29,30]. As a result, a single mRNA can be regulated by multiple miRNAs, while each miRNA can regulate numerous genes. miRNAs are indispensable in early embryonic development, influencing critical events such as brain morphogenesis and maternal mRNA clearance [31,32]. In the context of NC development, miRNAs dysregulation has been implicated in neurocristopathies, including DiGeorge syndrome and congenital central hypoventilation syndrome [25,33,34]. Furthermore, miRNAs are often dysregulated in tumours, acting as either oncogenic (*oncomiRs*) or tumour-suppressive (*anti-oncomiRs*) regulators [35,36]. NC-derived cancers, such as neuroblastomas and melanomas, share molecular features with developmental processes, highlighting the need for further exploration into the roles of miRNAs in both NC development and cancer progression [33].

In this study, we aim to identify and characterize miRNAs implicated in NC development through gain-of-function experiments in zebrafish embryos. By leveraging a double-transgenic zebrafish model (Tg(*sox10*:eGFP, *sox10*:mRFP)), we isolated NC and non-NC populations via fluorescence-activated cell sorting (FACS) and conducted transcriptomic analyses. Our findings reveal that some specific miRNAs regulate key genes involved in NC-derived structures, particularly craniofacial cartilage and melanophores through interactions with transcription factors such as Sox9b and Runx3. Notably, these miRNAs share regulatory mechanisms with tumour suppressor pathways, suggesting a broader role in cellular plasticity and differentiation. This research advances our understanding of vertebrate embryogenesis and provides valuable insights into the molecular mechanisms underlying neurocristopathies, NC-derived cancers, and tumorigenesis, underscoring the need for further investigation into the regulatory functions of miRNAs in both developmental and pathological contexts.

## 2 Results

### 2.1 miR-133a, miR-199-2 and miR-338-3p are overrepresented in NCC

Using the double transgenic zebrafish line Tg(*sox10*:eGFP,*sox10*:mRFP) [37], we separated the NCC and non-NCC population with FACS from 16 hpf (before the NCC migration and differentiation of NC derivatives) and 28 hpf embryos (after NCC migration and during differentiation of derivatives) and performed RNA-seq on each subpopulation.

At 16 hpf, we identified a total of 225 miRNAs, with 60 and 57 being miRNAs specific to NCC and non-NCC populations, respectively (Figure 1a; Fold Enrichment > 2, Supplementary Table ST2). In the NCC subpopulation, we found several previously reported NCC-associated miRNAs, such as miR-96 and miR-204 [38,39]. At 28 hpf, we identified 50 miRNAs in NCC, of which 12 miRNAs were overrepresented compared to non-NCC and one uniquely present in this subpopulation (miR-24, Figure 1b). Conversely, the non-NCC population at 28 hpf contained a total of 187 miRNAs, with 138 unique to this subpopulation and 17 overrepresented compared to NCC.

**Figure 1.**
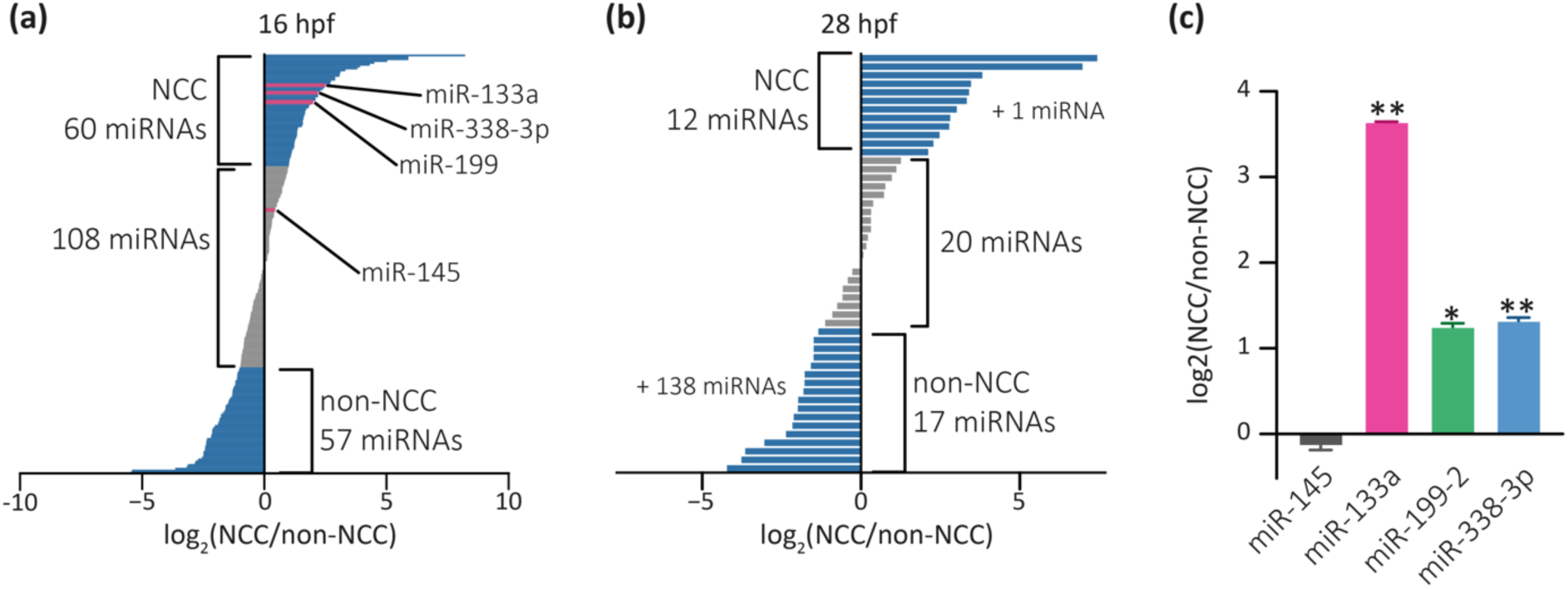
Identification of differentially expressed miRNAs in NCC and non-NCC from 16 hpf (a) and 28 hpf (b) transgenic zebrafish samples. Over-represented miRNAs in either cell population are shown in blue (log_2_(NCC/non-NCC) > 2). miR-145, miR-133a, miR-1SS-2 and miR-338-3p are highlighted in pink. (c) Quantification of miR-145, miR-133a, miR-1SS-2 and miR-338-3p levels in NCC by RT-qPCR, compared to non-NCC; levels are, expressed as log_2_(NCC/non-NCC). Statistical analysis was performed using a two-tailed Student’s t-test, *p ≤ 0.05, **p ≤ 0.01, n = 3, mean ± SEM.

Further analysis was performed on miRNAs over-represented in NCC at the 16 hpf stage in zebrafish, prior to NCC differentiation and migration. This early developmental time point was selected because it allows for the identification of miRNAs potentially involved in the initial specification and commitment of NCC, minimizing confounding effects from downstream lineage-specific expression. Based on bioinformatics analyses including STRING protein-protein interaction networks Gene Ontology (GO) classifications and KEGG pathway enrichments (Figure 2, Supplementary Figure S 1, 2, 3), we identified miR-133a, miR-199-2, and miR-338-3p as key candidates for further investigation. Besides, a literature review (see below) supported their potential roles in NCC specification and cancer biology.

**Figure 2.**
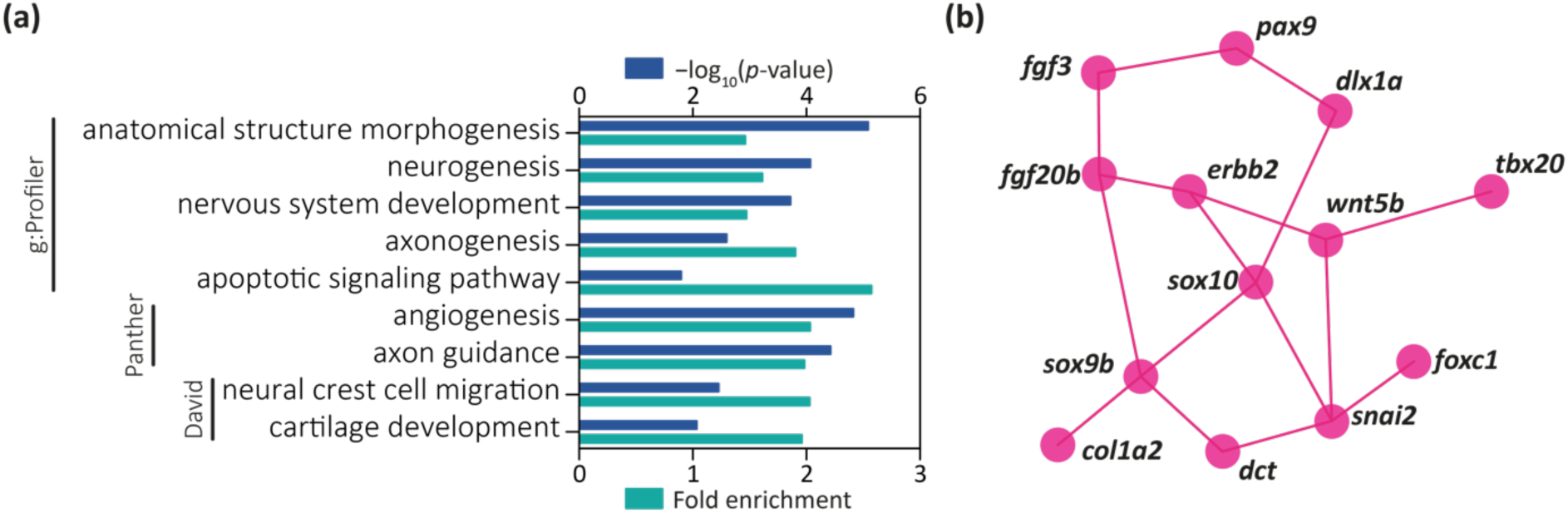
GO and STRING analysis of miR-133a putative target genes. Similar analyses for miR-1SS-2 and miR-338-3p are presented in Supplementary Figures S2 and S 3. (a) Selected overrepresented “Biological Processes” terms associated with miR-133a putative target genes, as identified by g:Profiler, Panther, and David. (b) Selected miR-133a putative target genes and their predicted interactions based on STRING. For the complete interaction network, refer to Supplementary Figure S 1.

Stem-loop RT-qPCR was performed on total RNA extracted from freshly dissociated, fluorescently sorted cells of Tg(sox10:eGFP,sox10:mRFP) [37] embryos at 16 hpf. miRNAs miR-133a, miR-199-2, and miR-338-3p were found to be overexpressed in NCC (Figure 1c), consistent with RNA-seq data. As a control, we analysed miR-145 expression, showing similar expression levels in NCC and non-NCC populations, as observed in the RNA-seq results (Figure 1a, c).

#### 2.1.1 miR-133a

miR-133a was initially identified in muscle tissue of zebrafish embryos at 48 hpf [40,41], and observed in the mouse heart and cerebral cortex [42]. It has been proposed as a biomarker for myocardial infarction due to decreased expression under this condition [43]. miR-133a suppresses the gene *Col1a1* in myocardial fibrosis [44] and aids in cardiac tissue regeneration [19,45]. In mice, miR-133a is co-transcribed with miR-1, both regulating processes in skeletal muscle development [46]. In cancer, miR-133a functions as a tumour suppressor by modulating FSCN1 expression in breast cancer [47] and, along with miR-145, in bladder and oesophageal cancers. Together with miR-1, it regulates proteins like TAGLN2 and PNP, inhibiting tumour proliferation in renal, bladder, prostate, and rhabdomyosarcoma cancers, and helping to prevent myocardial ischemia [48–52]. Circular RNAs circFUT10 and circ_nxc1 inhibit miR-133a, promoting mioblastomas and myocardial ischemia, respectively [53,54]. Although these studies were conducted in various model systems and pathological contexts, they collectively support a broader role for miR-133a in regulating cell cycle progression and cytoskeletal organization.

We retrieved a total of 2966 genes containing miR-133a putative binding sites from the TargetScan database (context score+ between-0.01 and-1.05, Supplementary Table ST3). According to g:Profiler (Padj < 0.05, g:SCS Threshold), multiple terms associated with neuronal development were overrepresented (Figure 2a, Supplementary Table ST4). Additional terms, such as "apoptosis signalling pathways," "anatomical structure development," and "developmental processes" were also identified, though neuronal development terms were the most predominant. Similar results were obtained using Panther (p-value < 0.05, FDR test), which also highlighted terms like "angiogenesis" and "axon development." DAVID (p-value < 0.05, FDR test) revealed NC-related terms, such as "neural crest cell migration" and "cartilage development." No NC development process pathway was overrepresented in this gene list (KEGG). Using the bioinformatic tool STRING, which generates interaction networks from gene lists, we identified central node genes that participate in multiple processes or interact with several genes within the list. This approach allowed for a focused analysis of miR-133a’s impact on genes that may have a significant influence on zebrafish embryonic development. By analysing a gene list with at least one putative miR-133a binding site and expression data from ZFIN for NCC or their derivatives, other central node genes were elucidated, including *ddx21*, *pax9*, *sox10*, *erbb2*, and *gf20b* (Figure 2b, Supplementary Figure S 1).

#### 2.1.2 miR-1SS-2

miR-199-2 has been identified in zebrafish development (starting from 20 hpf, primarily in the fins) as well as in various other vertebrates [41,55]. It has been shown to regulate ITGA3 expression (involved in cell adhesion), inhibiting cell migration and tumour invasion in squamous cell carcinoma [56]. In liver cancer cell cultures, miR-199-2 inhibits EMT [57] and regulates apoptosis in intervertebral disc cells by inhibiting MAP3K5 [58]. These findings position miR-199-2 as a tumour suppressor miRNA, like miR-133a. Its inhibitory effect on EMT in cancer may also be relevant to NC development and cell migration.

For GO and KEGG pathway analysis, we extracted a list of 3,000 genes with predicted binding sites for miR-199-2 from TargetScan, with context scores between −0.04 and −9.1, although a few overrepresented terms were found (Supplementary Tables ST5 and ST6). According to g:Profiler, 7 of 19 terms were related to nervous system development, while others were more general terms such as "multicellular organism development" and "developmental processes". Comparable results were obtained with Panther and DAVID, with terms like "axonogenesis" and "tissue development." A single KEGG pathway, "ErbB signalling pathway," was overrepresented. The absence of terms related to the NC, or its derivatives does not necessarily indicate that miR-199-2 is uninvolved in NC development, as it may regulate only a few key genes essential for this process. STRING analysis, using genes with putative binding sites for miR-199-2 and expression reported in ZFIN for NCC or their derivatives, identified several central node genes, including *foxd3*, *wnt5b*, *erbb2*, and various genes in the FGF signalling pathway (Supplementary Figure S 2).

#### 2.1.3 miR-338-3p

Bibliographic analysis revealed that miR-338-3p is linked to the nervous system, having been detected in neuron polyribosomes [59] and promoting nervous system development [60,61]. miR-338-3p inhibits EMT in gastric cancer by regulating ZEB2 [62], suppresses OSCC (oral cancer) cell growth by targeting NRP1 [63], and inhibits liver cancer cell invasion through SMO [64]. Like miR-133a and miR-199-2, miR-338-3p seems to be acting as a tumour suppressor by inhibiting cell proliferation.

For GO analysis, we generated a list of 3,001 genes with predicted miR-338-3p binding sites with TargetScan, yielding context scores between −0.04 and −1.48 (Supplementary Table ST7). g:Profiler identified only five biological process terms, three related to neuronal development and two to positive regulation of biological processes (Supplementary Table ST8). Panther identified additional terms like "embryonic development," while DAVID revealed terms such as "cartilage development," "fin morphogenesis," and "neural crest differentiation". No KEGG pathways were overrepresented. STRING analysis of genes with miR-338-3p binding sites and expression in NCC or derivatives identified *snai2*, *sox9b*, *kita*, and *pax2a* as central node genes (Supplementary Figure S 3).

These results suggest that miR-133a, miR-199-2, and miR-338-3p function as tumour suppressors, exhibiting reduced expression in various cancers. These miRNAs repress genes involved in EMT, migration, and cell proliferation—processes common to both cancer and NCC development. Although GO analysis revealed limited overrepresentation of NCC-specific terms, with only a few associated with miR-133a and miR-338-3p, there was a notable bias toward neuronal development. However, these miRNAs may still regulate key genes critical for NCC development, such as *soxE*, *snai1/2*, and *msx1/2/3*.

### 2.2 miR-133a, miR-199-2, and miR-338-3p are differentially expressed during zebrafish embryogenesis

Since the expression profiles of miR-133a, miR-199-2 and miR-338-3p in zebrafish remain incompletely characterized, we analysed their temporal expression from the 5 hpf stage onwards using RT-qPCR. Both miR-133a and miR-199-2 showed the lowest expression at expression at 5 hpf among the analysed stages (Figure 3a–b). Their expression begun to rise at 10 hpf, with a marked increase after 24 hpf. Notably, this upregulation at 10 hpf aligns with the expression patterns of NCC genes that contain binding sites for these miRNAs, such as *sox9b* and *foxd3 [65,66]*. Similarly, miR-338-3p was detected at 5 hpf—right after zygotic genome activation [67]—, but its levels decreased between 10 and 24 hpf before increasing again (Figure 3c). This pattern correlates with the downregulation of putative genes like *sox9b* and *snai1b [65,66]*. These findings suggest that these miRNAs exhibit temporally coordinated expression with their predicted target genes, highlighting their as potential role as regulators of gene expression during early zebrafish development.

**Figure 3.**
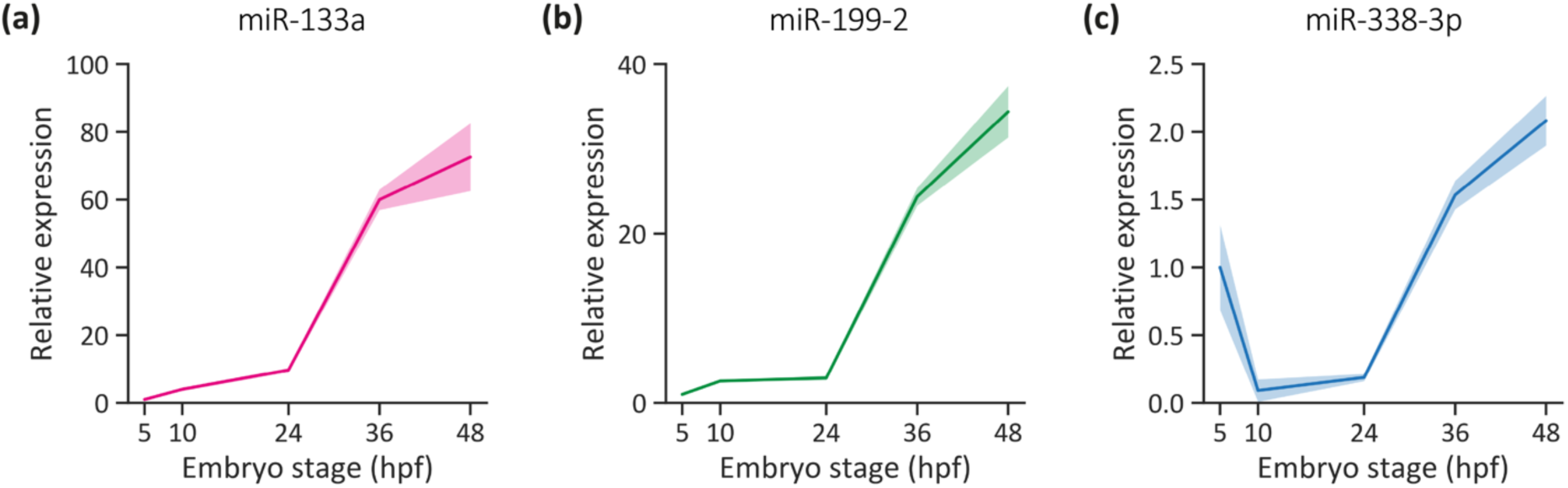
miRNA expression profiles during zebrafish embryonic development. (a-c) RT-qPCR quantification of miR-133a (a), miR-1SS-2 (b) and miR-338-3p (c) levels from 5 to 48 hpf. Normalized to 5 hpf stage, n = 3, mean ± SEM.

Given that miRNAs can simultaneously regulate multiple genes, and that each gene may be targeted by more than one miRNA, we filtered those genes present in at least two or all three predicted miRNAs putative target gene lists. Among these, only six NC-related genes were shared in all three lists, including *pdgIb*, *tbx20* and *ddx21* (Supplementary Table S9). Additionally, between 16 and 20 genes were common to two lists (Supplementary Table S9). Several of the identified target genes have been implicated in chondrocyte development, craniofacial structure formation, and melanophore differentiation. Given these associations, we decided to further investigate the roles of these miRNAs in these key biological processes.

We performed gain-of-function experiments in zebrafish embryos by microinjecting pri-miRNA constructs in tandem with a reporter dsRED sequence to increase the levels of miR-133a, miR-199-2, and miR-338-3p (dsRED-miR-133a, dsRED-miR-199-2, and dsRED-miR-338-3p, Supplementary Figure S 4a). As a control, embryos were injected with an mRNA generated from a construct containing only the dsRED sequence (dsRED-Control). We assessed during embryonic development both the presence of the reporter protein by fluorescence analysis and the pri-miRNA processing by stem-loop RT-qPCR. As expected, significant increase in miR-133a, miR-199-2, and miR-338-3p levels were observed in RNA samples from microinjected embryos collected at 24 hpf (Supplementary Figure S 4b). These results provided a foundation for further studies on the impact of miRNA overexpression on of NC derivative development.

### 2.3 miR-133a and miR-338-3p overexpression alters melanophore development

Zebrafish pigmentation is determined by three types of NC-derived chromatophores: melanophores (black pigment), iridophores (iridescent pigment that reflects light), and xanthophores (yellow pigment) [12]. In this study, we specifically examined the development of melanophores, which become visible around 25 hpf in the otic vesicle region. These cells subsequently expand across in the dorsal head region (Figure 4a) and along the embryonic lateral line at 48 hpf (Figure 4b) [12]. To investigate the role of miR-133a and miR-338-3p, both having putative binding sites to genes involved in melanophore development (*dct*, *kita*, *sox10*, *tyrp1a*) [14], we performed a quantitative analysis on melanophore number present in 48 hpf larvae following the overexpression of each miRNA. Overexpression of either miRNA led to a reduction in the melanophore number located in the dorsal head region (Figure 4c) and the lateral line (Figure 4d). Given that these miRNAs share putative target genes involved in melanophore development (like *dct*), we evaluated their potential synergistic effect through simultaneous overexpression. Co-overexpression of miR-133a and miR-338-3p produced a similar phenotype, suggesting that a maximal inhibitory effect on melanophore differentiation may already be achieved by either miRNA alone under the given experimental conditions. This aligns with the role of miRNAs as fine tuners of gene expression, where their regulatory impact does not necessarily scale linearly with increased expression. We further analysed the expression of *mi8a*, a master regulator of melanophore specification, and *dct* (dopachrome tautomerase), a key enzyme in melanin biosynthesis [14]. Notably, *mi8a* lacks predictive binding sites for either miRNA, and its expression remained unchanged following miR-133a or miR-338-3p overexpression (Figure 4e, Supplementary Figure S 5d).

**Figure 4.**
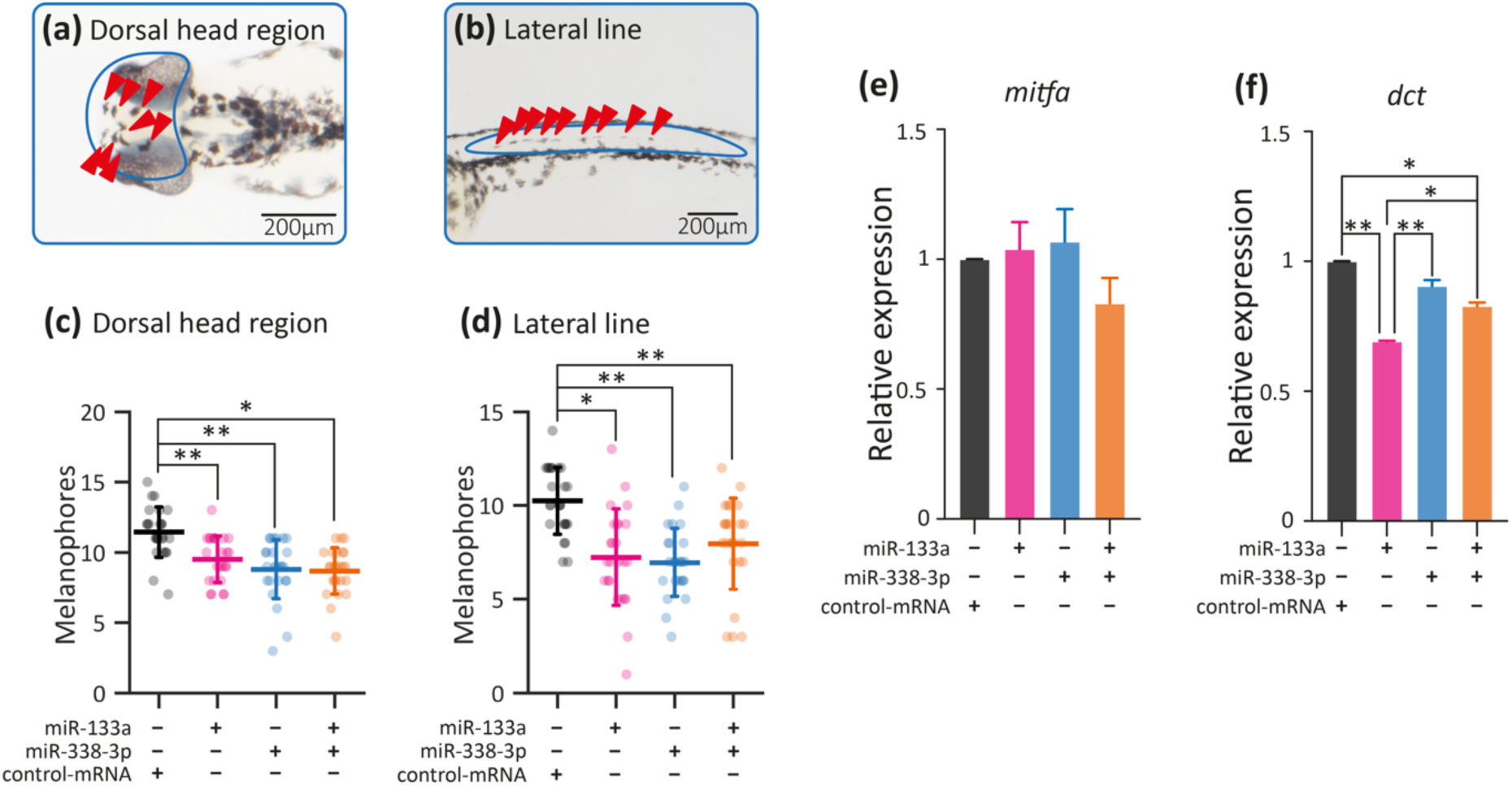
Melanophore development in miR-133a and miR-338-3p overexpressing embryos. (a-b) Representative images of melanophores in the head (a) and lateral line (b) regions. The analysed areas are highlighted in blue, with counted cells indicated by red arrowheads. Scale bars: 200 µm. (c-d) Quantification of melanophore numbers in the head (c) and lateral line (d) of 48 hpf larvae microinjected with dsRED-miR-133a, dsRED-miR-338-3p, both, or dsRED-control. Statistical analysis: ANOVA followed by Tukey’s post-hoc test; *p ≤ 0.01, **p ≤ 0.001, n = 25, mean ± SD. (e-f) mRNA expression levels of mitfa (e) and dct (f) in 24 hpf embryos microinjected with dsRED-miR-133a, dsRED-miR-338-3p, both, or dsRED-control. Statistical analysis: ANOVA followed by Tukey’s post-hoc test; *p ≤ 0.05, **p ≤ 0.01, n = 2, mean ± SEM.

In contrast, *dct* contains two predicted binding sites for miR-338-3p and multiple predicted sites for miR-133a (Supplementary Figure S 5e). Interestingly, while *dct* mRNA levels were unaffected in embryos overexpressing miR-338-3p, they were significantly reduced upon miR-133a overexpression (Figure 4f). This reduction may account for the observed decrease in pigmentation at 48 hpf in embryos microinjected with dsRED-miR-133a.

### 2.4 miR-133a and miR-338-3p overexpression adversely affect craniofacial cartilage development

Since all three miRNAs presented putative binding sites in genes related with chondrocyte development (*sox9b*, *runx3*, *col2a1b*, *col1a2*) [23,68,69], we designed gain-of-function experiments to evaluate cartilage formation. Embryos microinjected with dsRED-miRNA constructs were developed until 5 days post-fertilization (dpf), then fixed and stained with Alcian Blue (Figure 5a-b). Overexpression of miR-199-2 did not produce any detectable changes in cartilage morphogenesis within the analysed parameters (Figure 5d, h-i). In contrast, miR-133a overexpression led to a significant reduction in Meckel’s cartilage length and angle (ML and MAn, respectively; Figure 5c,f-g), as well as a decrease ceratohyal angle (ChAn) and an increased ceratohyal-to-fin distance (ChD) (Figure 5f-g). Similarly, larvae overexpressing miR-338-3p exhibited shorter palatoquadrate cartilages (PQ; Figure 5e, j), reduced Meckel’s cartilage angles, and an increased ceratohyal angle (ChAn; Figure 5k).

**Figure 5.**
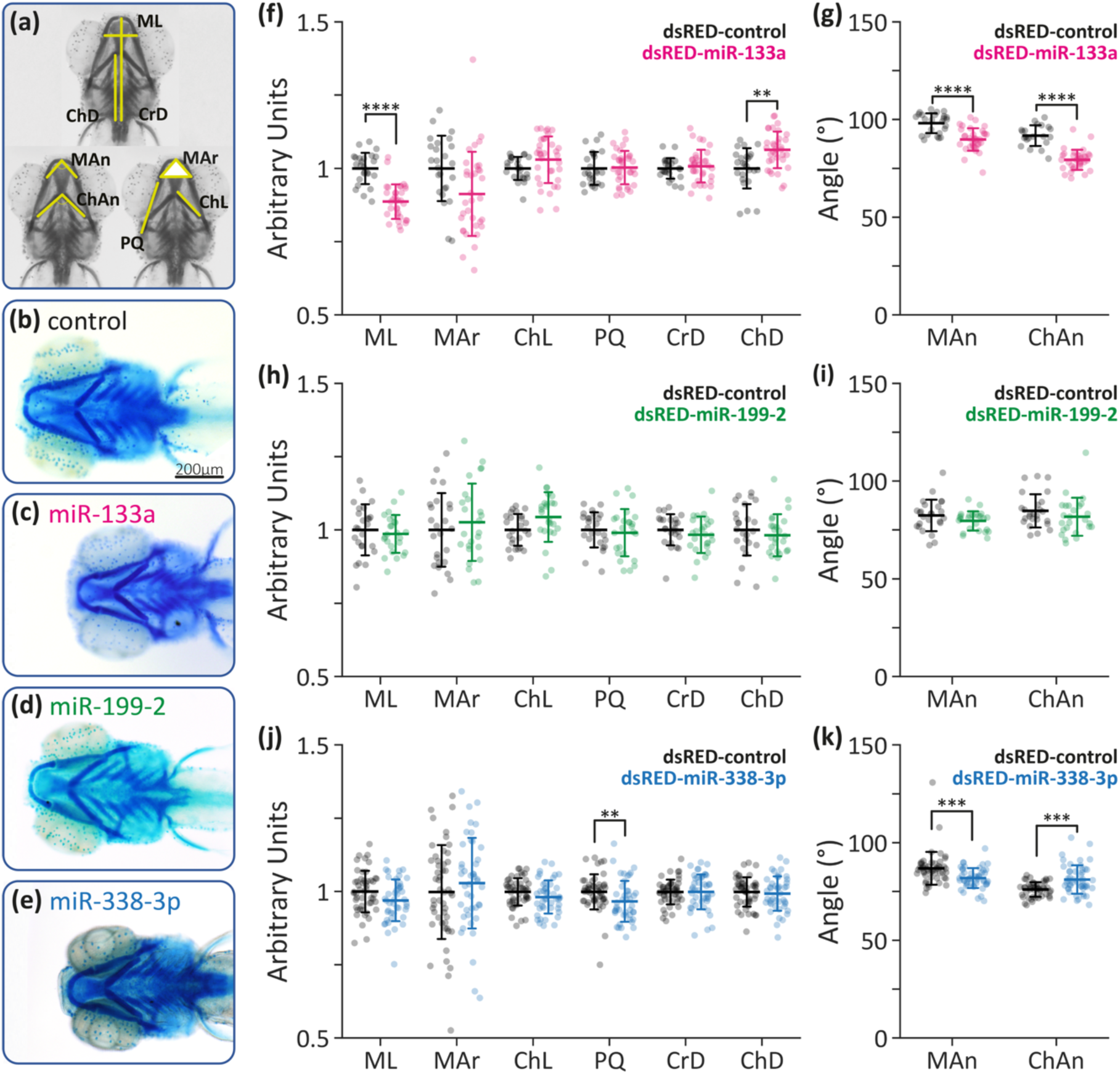
Craniofacial development of zebrafish specimens overexpressing miR-133a, miR-1SS-2 and miR-338-3p. (a) Schematic representation of the craniofacial parameters analysed: ML: Meckel’s cartilage length; MAr: Meckel’s cartilage area; ChL: Ceratohyal mean length; PQ: palatoquadrate mean length; CrD: cranial distance (from fins to Meckel’s cartilage midpoint); ChD: ceratohyal distance (from fins to ceratohyal cartilage joint); MAn: Meckel’s cartilage angle; ChAn: ceratohyal angle. (b-e) Representative ventral view images of Alcian blue-stained cartilage in 5-dpf larva, with the head positioned to the left. Images show specimens developed from embryos microinjected with: dsRED-control (b), dsRED-miR-133a (c), dsRED-miR-1SS-2 (d) and dsRED-miR-338-3p (e). (f-k) Craniofacial measurements larvae microinjected with: dsRED-miR-133a (f-g), dsRED-miR-1SS-2 (h-i), and dsRED-miR-338-3p (j-k). Statistical analysis; Two-tailed Student’s t-test; *p ≤ 0,05, **p ≤ 0,01, ***p ≤n0,001, ****p ≤ 0,0001, n = 25 for miR-1SS-2, n = 22 for miR-133a, n = 48 for miR-338-3p, mean ± SD. Scale bar in (b): 200 µm.

Since miR-133a and miR-338-3p are predicted to co-target several genes associated with craniofacial development (like *sox9b*, *sox4a* and *wnt9b*) [21,70,71], we evaluated their potential synergistic effect. Our results revealed a significant reduction in both the Meckeĺs and ceratohyal angles, consistent with the trends observed when each miRNA was overexpressed individually (Figure 6a-b). Additionally, co-microinjected larvae exhibited an increase in the Meckel’s cartilage area (Figure 6a-b). Interestingly, while the individual overexpression of dsRED-miR-133a or miR-338-3p led to reduced lengths of the Meckel (ML) and palatoquadrate (PQ) cartilages, and the distance from the fins to the ceratohyal cartilages (ChD), their dual overexpression restored these parameters to control levels. This suggests that the simultaneous presence of miR-133a and miR-338-3p, particularly in NC tissues where they are not typically co-expressed, may exert a complex regulatory influence on craniofacial development by modulating shared target genes. Alternatively, the restoration of ML values upon co-overexpression may indicate that each miRNA regulates distinct sets of genes with opposing effects on this parameter, ultimately balancing one another’s impact. Further investigation is required to elucidate the precise molecular mechanisms underlying this interaction.

**Figure 6.**
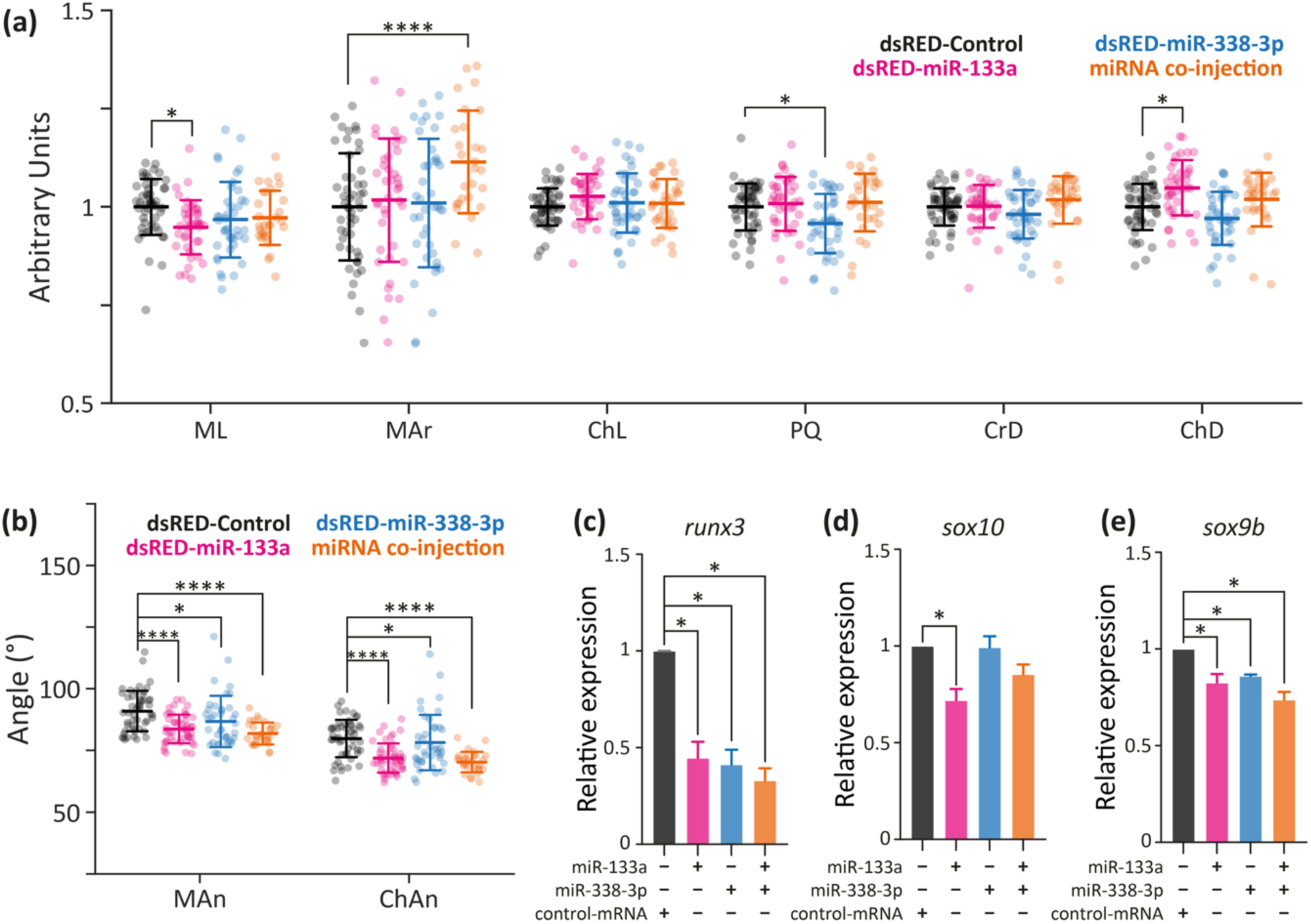
Synergistic effect of miR-133a and miR-338-3p on craniofacial development. (a-b) Craniofacial measurements in larvae microinjected with dsRED-miR-133a, dsRED-miR-338-3p, both, or dsRED-control. Statistical analysis: ANOVA followed by Tukey’s post-hoc test; *p ≤ 0.05, ****p ≤ 0.0001, n = 30, mean ± SD. (c-e) mRNA expression levels of runx3 (c), sox10 (d) and sox9b (e) in 24 hpf embryos microinjected with dsRED-miR-133a, dsRED-miR-338-3p, both, or dsRED-control. Statistical analysis: ANOVA followed by Tukey’s post-hoc test; *p ≤ 0.05, n = 2, mean ± SEM.

To further elucidate the molecular mechanisms underlying the observed craniofacial defects, we quantified the mRNA expression levels of *runx3*, *sox10*, and *sox9b*—key regulators of cartilage development [23]—using RT-qPCR. This analysis revealed a significant reduction in *runx3* expression in embryos microinjected with dsRED-miR-133a and dsRED-miR-338-3p, despite *runx3* containing only a single putative binding site for miR-133a (Figure 6c, Supplementary Figure S 5c). The effect of miR-338-3p on *runx3* expression may be indirect, potentially resulting from the dysregulation of upstream genes within the gene regulatory network (GRN) or a reduction in the specific cell population expressing *runx3*. Regarding *sox10*, mRNA levels were specifically reduced in dsRED-miR-133a microinjected embryos, consistent with the presence of a predicted binding site for this miRNA in the 3’UTR of *sox10* (Figure 6d, Supplementary Figure S 5a). Given that *sox10* is an early NC development marker [9], its downregulation could contribute to the observed craniofacial abnormalities. Additionally, overexpression of both miR-133a and miR-338-3p resulted in decreased *sox9b* mRNA levels, likely due to the presence of multiple binding sites for these miRNAs in its 3’UTR (Figure 6e, Supplementary Figure S 5b). Since *sox9b* is a crucial regulator of chondrogenesis and NC-derived cartilage formation, its downregulation could be a key factor underlying the structural defects observed.

### 2.5 miR-133a and miR-338-3p bind to sox9b 3’UTR

The observed reduction in *sox10*, *sox9b*, *dct*, and *runx3* mRNA levels following the overexpression of miR-133a and miR-338-3p may be attributed to multiple mechanisms. These include a decrease in the specific cell populations expressing these genes, a reduction in the availability of transcription factors that activate their expression, or a direct interaction between the miRNA and the 3’UTR of the target gene. Such interactions can lead not only to mRNA degradation but also to translational repression [26].

To determine whether these mRNA reductions stem from direct regulation by miR-133a or miR-338-3p, we performed a reporter assay using the *d4GFPn* gene to assess *sox9b* translational repression. *Sox9b* was selected due to its critical role in cartilage development and the presence of multiple predictive miRNA binding sites in its 3’UTR. Briefly, we used different constructs in which the d4GFPn coding sequence fused to specific fragments of the *sox9b* 3’UTR. The first construct, *sox9b*-3’UTR-1, contains a single putative binding site for miR-133a and another for miR-338-3p, while the second construct, *sox9b*-3’UTR-2, harbours two putative binding sites exclusively for miR-338-3p (Figure 7g). The corresponding mRNAs were transcribed *in vitro* and co-injected into 1-cell stage embryos alongside dsRED-miR-133a, dsRED-miR-338-3p, both, or dsRED-control (Supplementary Figure S 5). Embryos were subsequently imaged at the 50%-epiboly stage, and GFP fluorescence was quantified using QuantiFish. As an additional control, a third construct (control-3’UTR) was designed, in which d*4GFPn* was fused to the SV40-poly(A) sequence, ensuring that any observed effects were specifically attributed to *sox9b* 3’UTR sequences.

**Figure 7.**
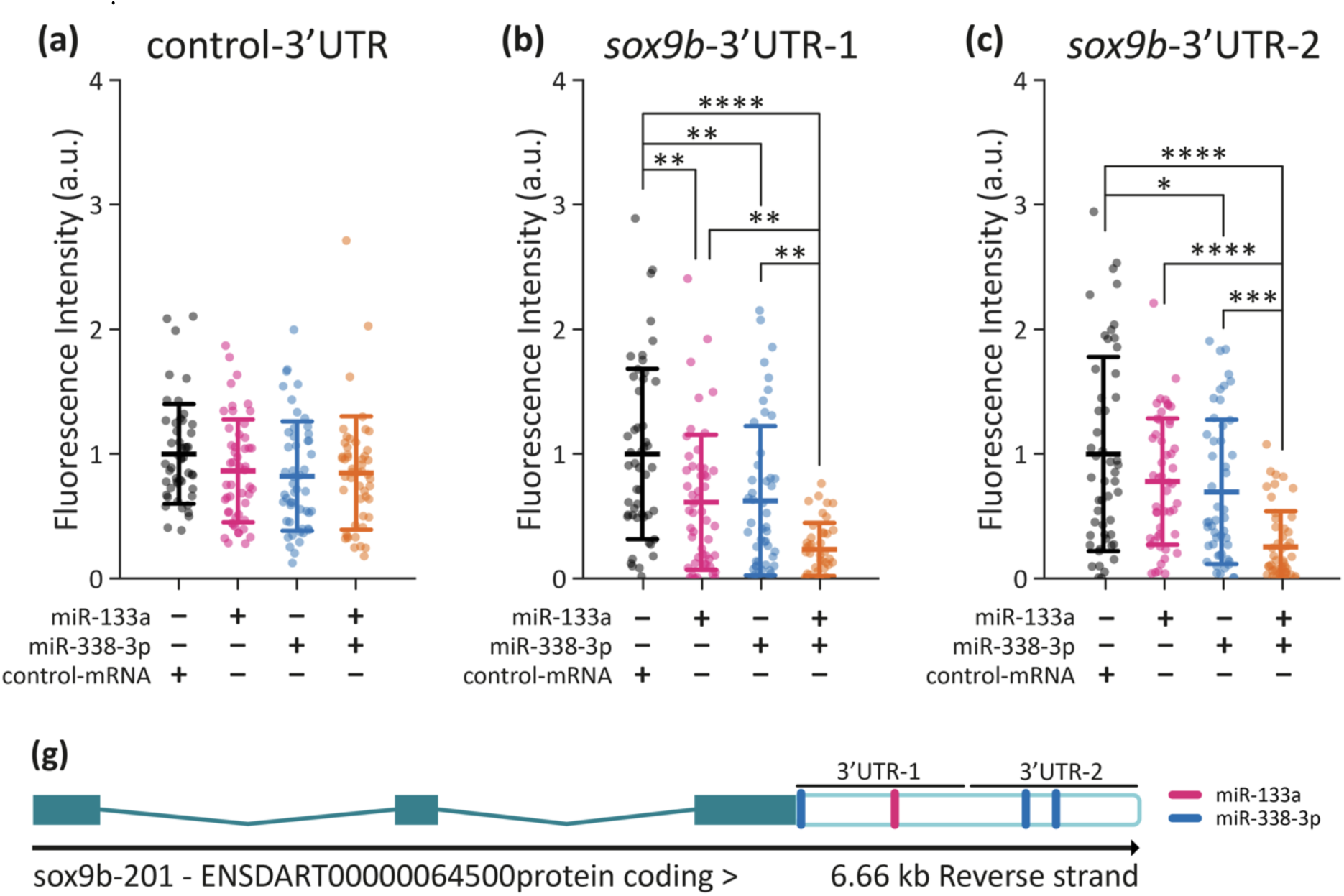
Interaction between miR-133a, miR-338-3p, and the soxSb 3’UTR. (a-c) Quantification of d4GFPn ffuorescence in embryos co-injected with either the control-3’UTR, soxSb-3’UTR-1, or soxSb-3’UTR-2, alongside dsRED-miR-133a, dsRED-miR-338-3p, both, or dsRED-control. Statistical analysis was performed using ANOVA followed by Tukey’s post-hoc test; *p ≤ 0.05, **p ≤ 0.01, ***p ≤ 0.001, ****p ≤ 0.0001, n = 45, mean ± SD. (g) Schematic representation of the soxSb gene structure (green), highlighting the 3’UTR (empty box) and the predicted binding sites for miR-133a (blue) and miR-338-3p (pink). The soxSb-3’UTR-1 and soxSb-3’UTR-2 fragments, used in the d4GFPn reporter constructs are indicated.

In embryos microinjected with *sox9b*-3’UTR-1, overexpression of both miR-133a and miR-338-3p significantly suppressed *d4GFPn* translation, with the strongest effect observed upon co-microinjection of both miRNAs (Figure 7b). Conversely, embryos injected with *sox9b*-3’UTR-2, exhibited a significant reduction in fluorescence only when dsRED-miR-338-3p was present (Figure 7c). No significant changes were detected in embryos microinjected with the control-3’UTR, regardless of miRNA treatment (Figure 7a). These findings align with RT-qPCR results, reinforcing the notion that miR-133a and miR-338-3p directly regulate *sox9b* expression by targeting its 3’UTR. This regulation likely occurs through translational repression, mRNA degradation, or a combination of both mechanisms. The observed reduction in *sox9b* mRNA levels in RT-qPCR analyses may, at least in part, result from miRNA-induced mRNA destabilisation.

### 2.6 miR-338-3p overexpression results in higher NCC number

The effects described thus far could stem from impaired cell differentiation, disruptions in migration patterns, and/or dysregulated NCC proliferation. To quantitatively evaluate changes in the NCC population, we microinjected Tg(*sox10:*GFP) embryos with dsRED-miR-133a, dsRED-miR-338-3p, or a dsRED-control, followed by flow cytometry analysis to quantify GFP-positive cells (Figure 8a). While miR-133a overexpression did not significantly alter NCC numbers, miR-338-3p overexpression resulted in a notable increase in NCC abundance (Figure 8b). Interestingly, the co-expression of both miRNAs restored NCC numbers to control levels, suggesting a potential compensatory mechanism. The proliferation and differentiation of NCC are tightly regulated by numerous transcription factors and signalling pathways [72]. Although miR-133a is predicted to target genes associated with cell proliferation, its overexpression did not significantly impact NCC numbers at 18 hpf. In contrast, miR-338-3p overexpression resulted in a marked increase in NCC abundance. This effect may stem from miR-338-3p’s regulatory influence on genes involved in cell proliferation, such as *wif1* and *ddx21*, both of which have been implicated in NC expansion and developmental patterning [73–76]. The increase in NCC following dsRED-miR-338-3p overexpression may contribute to the defects in melanophore and chondrocyte development described above. Given miR-338-3p’s involvement in both early (proliferation) and later (differentiation) stages of NC development, its dysregulation likely disrupts the balance between cell number and lineage specification, ultimately leading to the observed phenotypic abnormalities. In contrast, since miR-133a overexpression did not affect NCC proliferation, the associated defects in melanocyte and craniofacial development are more likely due to miR-133a-mediated regulation of genes involved in NCC specification and differentiation, rather than alterations in cell number.

**Figure 8.**
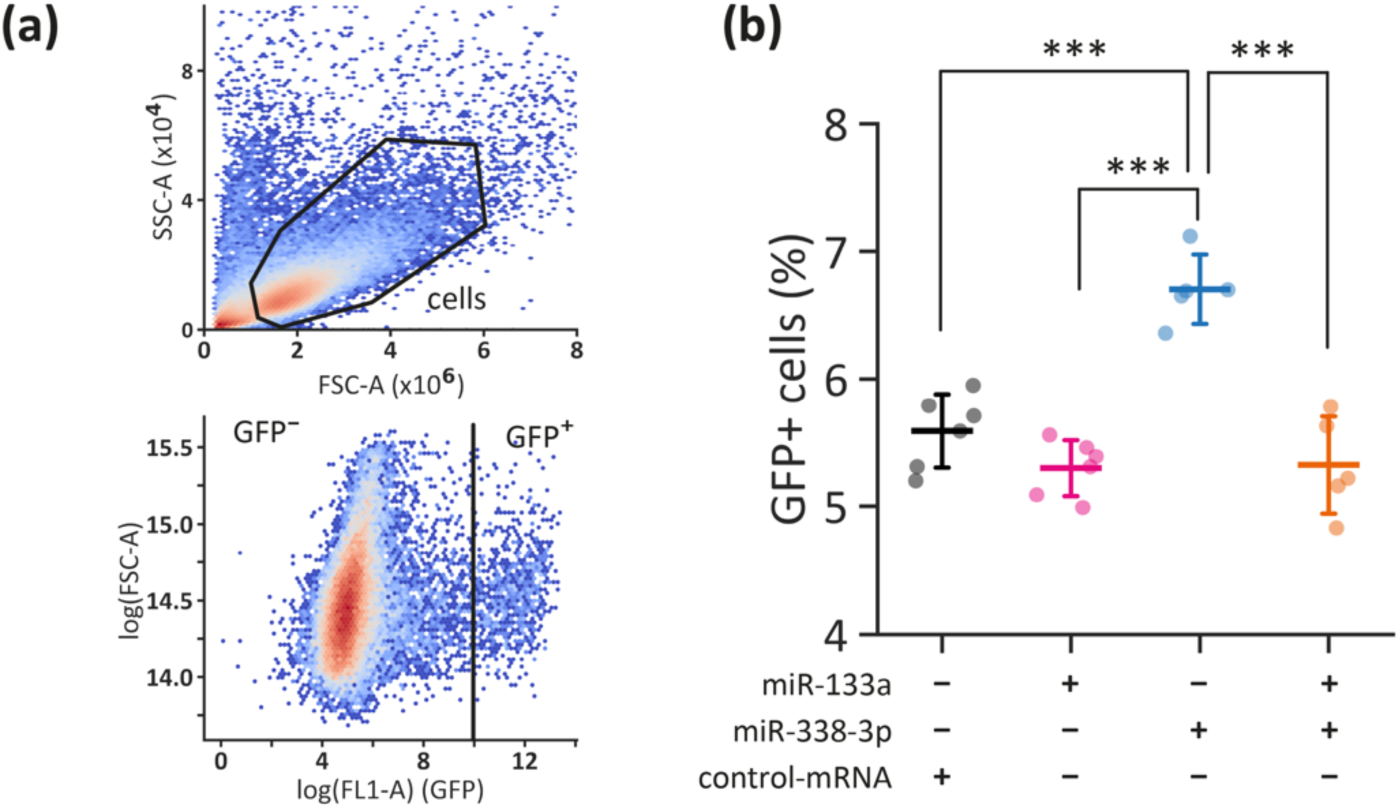
NCC proliferation. (a) Gating strategy used to identify zebrafish cells [77] (upper panel) and distinguish GFP-negative and GFP-positive cell populations (lower panel). (b) Flow Cytometry-based quantification of NCC proportions in 18 hpf embryos microinjected with dsRED-miR-133a, dsRED-miR-338-3p, both, or dsRED-control. Statistical analysis: ANOVA followed by post-hoc Tukey test, ***p ≤ 0.001, n = 5, mean ± SD.

## 3 Discussion

MiRNAs are pivotal regulators of embryonic development and numerous diseases, including cancer, where they fine tune gene expression at multiple levels [24,78]. Due to their ability to target multiple mRNAs simultaneously [79] and considering the extensive network of transcription factors, enzymes, and structural proteins involved in the GRN [6,80] of embryonic development- it is likely that many miRNA-mRNA interactions remain undiscovered. The parallels between NC development and cancer biology, particularly in processes such as EMT, migration, and invasion, suggest a shared regulatory landscape. Insights gained from one context can, therefore, enhance our understanding of the other.

Building on this concept, we selected miRNAs with established roles in tumorigenesis to investigate their regulatory influence on NC-derived tissues. Specifically, the roles of miR-133a, miR-199-2, and miR-338-3p, that are highly expressed in the premigratory NCC of *Danio rerio.* These miRNAs, which are conserved exclusively among vertebrates, have been implicated in cancer-related processes such as EMT, cytoskeletal reorganization, proliferation, migration, and apoptosis, as supported by both literature and bioinformatic analyses. Through this study, their contributions to NC-derived structures pathways were explored.

The literature suggests that miR-199-2 plays a role regulating EMT and tumour cell migration, processes that also crucial during early NC development [57,58,81]. However, no direct evidence has yet linked miR-199-2 to the regulation of specific NC derivatives. Notably, putative target genes of miR-199-2 include key NC specifiers such as *foxd3 [6]*, as well as transcription factors involved in signalling pathways essential for NC specification, including Wnt and FGFs [82]. While miR-199-2 overexpression did not result in detectable craniofacial defects, its role in NC development cannot be ruled out, particularly in processes such as cell migration and iridophore differentiation. This possibility is supported by the presence of putative binding sites in genes implicated in these processes, such as *ltk [16]*, as well as its reported involvement in tumorigenesis and cell adhesion. Further studies, including functional assays and lineage tracing, are necessary to elucidate the precise role of miR-199-2 in NC development and its potential contribution to these biological processes.

The proper formation of craniofacial structures relies on the coordinated proliferation and positioning of cartilage progenitor cells [21]. Transcription factors governing craniofacial development have been extensively characterized in *Danio rerio*, mice, and cultured cells. Our findings indicate that miR-133a and miR-338-3p regulate the expression of *sox9b*, a critical transcription factor required not only for craniofacial cartilage formation but also for melanophore differentiation. Here, we have demonstrated that these miRNAs repress *sox9b* expression at both the transcriptional and translational levels by directly interacting with its 3ʹUTR. Notably, co-overexpression of miR-133a and miR-338-3p exacerbates craniofacial malformations, suggesting a synergistic effect on cartilage development. This outcome likely results from their combined modulation of upstream regulators of chondrogenesis, highlighting their broader impact on NC-derived structures.

We also found that miR-133a and miR-338-3p regulate *runx3*, a crucial gene expressed in the pharyngeal endoderm that is essential for the activation of downstream cartilage GRN [. The reduction in *runx3* expression following miR-133a and miR-338-3p overexpression mimics phenotypes observed in *runx3* morphants, including defects in viscerocranial and anterior neurocranial cartilage [23]. While miR-133a likely directly targets *runx3*, miR-338-3p appears to act indirectly, potentially through the modulation of other transcription factors involved in chondrogenesis. These findings highlight the complexity of miRNA-mediated regulation, which can influence developmental pathways both directly and indirectly through highly interconnected GRNs.

In melanogenesis, both miR-133a and miR-338-3p also play significant regulatory roles. Vertebrate melanocytes (melanophores in fish) are critical for pigmentation, mating behaviours, individual recognition, and UV protection. They are also implicated in various pigmentation disorders, including albinism, vitiligo, and melanoma [84]. Our data indicate that miR-133a reduces *dct* expression, which may account for the observed decrease in melanocyte numbers. Interestingly, although *sox10* is required for the activation of *mi8a [14]*, we did not detect a corresponding reduction in *mi8a* expression. This aligns with reports that zebrafish *mi8a* mutants retain a small population of pigmented cells that depend on *sox9b [14]*. Since *sox9b* is transiently required in melanocyte precursors, the downregulation of *sox9b* by miR-133a and miR-338-3p may disrupt early melanocyte activation, ultimately leading to reduced melanogenesis. The absence of detectable changes in the expression levels of *dct*—an enzyme required for melanin biosynthesis [14]—following miR-338-3p overexpression does not necessarily exclude miRNA-mediated regulation. It is possible that miR-338-3p modulates *dct* expression at the translational level rather than by inducing mRNA degradation, a mechanism consistent with the well-established dual role of miRNAs in post-transcriptional regulation. Further studies are needed to elucidate the precise molecular pathways through which these miRNAs influence zebrafish melanophore differentiation and pigmentation.

Contrary to its well-documented tumour-suppressor role [62–64], miR-338-3p overexpression led to an increased population of NC progenitor cells in zebrafish embryos. This unexpected outcome likely reflects the widespread effects of ubiquitous miRNA overexpression, which impacts all embryonic cells rather than being confined to NC tissues. The elevated proliferation of progenitor cells may contribute to developmental defects observed at later stages, highlighting the need for more refined spatial and temporal analyses to delineate the specific roles of miRNAs during development.

While GO analyses did not reveal extensive enrichment of NC-related pathways, our findings strongly suggest that miR-133a and miR-338-3p play crucial roles in NCC development. These miRNAs likely exert their effects through both direct and indirect regulatory mechanisms, influencing key transcription factors such as *sox9b* and *runx3*. The interplay between these miRNAs and their target genes underscores their broader significance in cellular plasticity and differentiation.

It is important to note that bioinformatic predictions of miRNA-mRNA interactions often lack experimental validation. Demonstrating the co-expression of miRNAs and their targets within the same cell type is essential for confirming functional interactions [85–88]. Future studies employing *in situ* hybridization will be valuable for validating the co-localization of miRNAs and their putative targets, such as *sox10*, *runx3*, and *dct*. The intricate overlap of GRNs regulating various NC derivatives suggests that miRNA overexpression likely affects multiple tissues simultaneously. For instance, the dual regulatory effect of miR-133a and miR-338-3p on *sox9b* influences both craniofacial cartilage formation and melanogenesis, demonstrating how miRNAs orchestrate broad developmental processes.

In conclusion, our findings highlight the essential role of precise miR-133a and miR-338-3p regulation in the development of NC-derived structures in *Danio rerio*, particularly craniofacial cartilage and melanophores. The craniofacial abnormalities observed in miRNA-overexpressing embryos likely stem from the direct repression of *sox9b* expression, while the reduction in the number of melanophore may result from downregulation of *dct*. Beyond their developmental roles, the shared regulatory features between NC formation and tumorigenesis underscore a broader function for these miRNAs in modulating cellular plasticity, proliferation, and differentiation. These insights not only deepen our understanding of miRNA-mediated control during vertebrate embryogenesis but also highlight the potential of miR-133a and miR-338-3p as therapeutic targets in neurocristopathies and cancer. Future studies should further investigate their mechanistic roles and translational relevance in both developmental and pathological contexts.

## 4 Materials & Methods

### 4.1 Zebrafish care

All zebrafish were handled in accordance with national and international guidelines [89]. All protocols involving zebrafish from the Calcaterra Lab were approved in advance by the *Comité de Bioética para el Manejo y Uso de Animales de Laboratorio* of the Facultad de Ciencias Bioquímicas y Farmacéuticas, UNR (files 6060/374, resolution 207/2018). Adult zebrafish were maintained at a constant temperature of 27 ± 1 °C under a 14:10 h light/dark cycle. [89] Wild-type (WT) fish from the AB strain (ZIRC, Oregon University, OR, USA) were used in this study. The transgenic lines used included Tg(*actb1*:eGFP), Tg(*sox10*:eGFP, *sox10*:mRFP) [37] and Tg(*sox10*:eGFP).

Zebrafish embryos were obtained according to the methodology described by Kimmel *et al [90].* Briefly, selected adults were kept overnight at 28.5 °C in breeding tanks, with males and females separated in a 3-to-4 ratio, respectively. Embryos were staged by visualization under stereoscopic microscope (Olympus MVX10 stereoscopic microscope and Olympus C-60 ZOOM digital camera) based on morphological development in hours or days post-fertilization (hpf or dpf) at 28 °C. [90]

### 4.2 Flow cytometry and FACS

Embryos at 16 and 28 hpf were manually dechorionated and deyolked using Ginsburg buffer (111.22 mM NaCl, 3.35 mM KCl, 2.7 mM CaCl₂, 2.38 mM NaHCO₃) through gentle pipetting and vortexing (600 rpm, 5 min). The samples were then centrifuged at 2500 rcf for 3 min at 4 °C, resuspended in 1X PBS containing 0.125% (w/v) Trypsin, and incubated at 37 °C for 15 min to facilitate dissociation. Following incubation, the samples were centrifuged (500 rcf, 15 min, 4 °C), resuspended in 500 µL 1X PBS, filtered through a 40 µm mesh, and counted.

To calibrate fluorescence-based cell sorting, wild-type and Tg(*actb1*:eGFP) lines were used as controls, representing 0% and 100% eGFP+ cells, respectively. For Fluorescence-Activated Cell Sorting (FACS) and Flow Cytometry cell counting, the transgenic lines Tg(*sox10*:eGFP, *sox10*:mRFP) and Tg(*sox10*:eGFP) were employed., respectively. FACS was conducted using BD FACSAria II and FACSAria III systems (BD Biosciences, NJ, USA) at a flow rate of 30 µL/min. GFP+ cells were used to establish gating parameters for zebrafish cells [77]s., while mRFP fluorescence served as a control, achieving >99% mRFP+ within the eGFP+ population. Cells sorted by FACS were directly collected in TRIzol (Invitrogen) for subsequent RNA extraction.

Flow Cytometry cell counting was conducted using a BD Accuri C6 Plus system (BD Biosciences), with a flow rate of 30 µL/min, ensuring a minimum of 100,000 events were recorded per condition. Each condition was analysed in triplicate for each biological replicate. Data processing and analysis were performed using FlowJo software (BD Biosciences).

### 4.3 RNA-seq

Total RNA was extracted using TRIzol Reagent (Life Technologies) with DNase treatment to remove genomic contamination. RNA concentration and purity were assessed using the Qubit RNA BR Assay kit (Thermo Fisher Scientific), while RNA integrity was evaluated with the RNA 6000 Nano Bioanalyser 2100 Assay (Agilent). After confirming RNA quality, samples from different experimental groups were used for library construction at each developmental stage.

Small RNA libraries were prepared using the TruSeq Small RNA Library Prep kit (Illumina) according to the manufacturer’s protocol. Sequencing was performed on an Illumina NextSeq High Output platform in paired-end mode, generating 75 bp reads. Image analysis, base calling, and quality scoring were carried out using Illumina’s Real-Time Analysis software (v3.4.4). The cleaned-up reads were checked using FASTQC and then aligned to the zebrafish genome using Bowtie [91]. Aligned reads were counted using HTSeq [92]. Read counts for each sample included in the study were normalized after which the miRNA differential expression was analysed using DESeq2 [93].

### 4.4 Bioinformatics

Zebrafish (*Danio rerio*) pre-miRNA sequences were retrieved from Ensembl (Release: 89, Assembly: GRCz11) and miRBase (Version: 22.1). Putative miRNA target genes were identified using TargetScan Fish (targetscan.org/fish_62) [94]. Gene expression data for zebrafish were extracted from the EMBL-EBI Expression Atlas, based on experimental data from White *et al.,* 2017 [65,66].

Gene Ontology (GO) and KEGG metabolic pathways analyses were conducted using multiple bioinformatic tools: DAVID (Version: v2022q4; GO and KEGG, FDR and p-value < 0.05) [95], PANTHER (Version: 10.5281/zenodo.4495804; GO, FDR and p-value < 0.05) [96], and g:Profiler (Version: e111_eg58_p18_f463989d; GO and KEGG, g:SCS Threshold and *p*adj < 0.05) [97].

Statistical analyses were performed using Prism 9.5 (GraphPad Software, Boston, MA, USA). Depending on the experimental design, either a two-tailed t-test or ordinary/nested ANOVA followed by *post-hoc* Tukey’s test was applied. Statistical significance was set at p ≤ 0.05. Data visualization was performed with Prism 9.5 or with custom scripts using Python (v3.10), generated with pandas, seaborn and matplotlib libraries.

### 4.5 Reverse Transcription Followed by Quantitative PCR (RT-qPCR)

For RNA extraction, 35-45 embryos per condition were collected at the required developmental stage and rapidly flash-frozen in liquid nitrogen. Samples were either processed immediately or stored at -80 °C for long-term preservation. Embryos were homogenized in TRIzol, followed by chloroform extraction and isopropanol precipitation. Total RNA concentration was determined by measuring absorbance at 260 nm using a NanoVue spectrophotometer (GE Healthcare, Chicago, IL, USA).

Reverse transcription was performed using M-MLV reverse transcriptase (Promega, Madison, WI, USA) with 1 µg of RNA. Oligo-dT primers were used for mRNA reverse transcription, while specific stem-loop [98] oligonucleotides were designed for each miRNA. Quantitative PCR (qPCR) was conducted using HOT FIREPol EvaGreen qPCR Mix Plus (Solis Biodyne, Tartu, Estonia) on a RealPlex4 thermocycler (Eppendorf, Hamburg, Germany). Zebrafish *rpl13* and *eef1a1l1* cDNAs were amplified as internal references. Primer sequences are provided in Supplementary Table S6. Data analysis was carried out using REST2009 software (Qiagen, Hilden, Germany) [99], following MIQE guidelines [100] to ensure experimental accuracy and reproducibility.

### 4.6 miRNA overexpression

For the overexpression experiments, the genomic regions encoding zebrafish *miR-133a* (miRBase Accession: MIMAT0001830; chr2: 4,113,889–4,114,465), *miR-199-2* (miRBase Accession: MIMAT0003155; chr5: 1,376,464–1,377,047), and *miR-338-3p* (miRBase Accession: MIMAT0048673; chr3: 51,907,303–51,907,638) were PCR-amplified and cloned into the pSP64T-dsRED expression vector using *Eco*RI and *Xho*I restriction sites (primer sequences provided in Supplementary Table S1, with restriction sites in lowercase).

For mRNA synthesis, plasmids were linearized with either *Xba*I or *Bam*HI (Invitrogen, Carlsbad, CA, USA) and transcribed using the mMESSAGE mMACHINE® SP6 transcription kit (Invitrogen). A ds-RED control vector lacking miRNA sequences was prepared as a negative control.

Microinjections were performed at the one-cell stage, targeting the yolk just beneath the blastomere, using a gas-driven microinjection system (MPPI-2 Pressure Injector, Applied Scientific Instrumentation, Eugene, OR, USA). Each embryo received 1.25 ng of transcript and was maintained at 28 °C until the required developmental stage for analysis.

### 4.7 Reporter Assay

The *sox9b* d4EGFPn-3’UTR reporter constructs were originally generated by Steeman *et al [101]*. An empty pSP64-T-d4EGFPn vector was used as a control. Transcripts were microinjected into one-cell-stage embryos at a concentration of 1.5 ng, along with 1.25 ng of either miR-133a-dsRED, miR-199-2-dsRED, miR-338-3p-dsRED, or control-dsRED transcripts.

Fluorescence from D4EGFPn was assessed at the 50%-epiboly stage. At least 40 embryos per condition were imaged using an Olympus MVX10 stereomicroscope equipped with an Olympus C-60 ZOOM digital camera (Olympus, Tokio, Japan). Fluorescence intensity was quantified using QuantiFish software [102].

### 4.8 Alcian Blue Staining

To assess the effects on cranial structures, 30 larvae at 5 dpf were fixed overnight at 4 °C in 4% (w/v) paraformaldehyde prepared in 1X PBT (1X PBS with 0.1% (v/v) Tween-20). Following fixation, samples were washed four times with 1X PBT and stained according to previously established protocols [103].

Images were captured using an Olympus MVX10 stereomicroscope equipped with an Olympus C-60 ZOOM digital camera. Cranial cartilage parameters were measured using ImageJ software (National Institute of Health, Bethesda, MD, USA) [104] following previously reported methods [4].

### 4.9 Melanophore counting

Larvae at 48 and 72 hpf were anesthetized using 0.15 mg/mL Tricaine (ethyl 3-aminobenzoate methanesulfonate salt, Sigma-Aldrich, Ref. A5040) and positioned in Petri dishes containing 3% (w/v) methylcellulose. Using two 30 G needles, larvae were carefully oriented to obtain lateral and dorsal images under an Olympus MVX10 stereomicroscope equipped with an Olympus C-60 ZOOM digital camera. Imaging was performed using top illumination against a white background. Pigmented cell counts were conducted manually by analysing the acquiring images with Fiji software.

## Abbreviations

miRNA: microRNA
NC: neural crest
GRN: gene regulatory network
dpf: days post-fecundation
hpf: hours post-fecundation

## 5 CRediT authorship contribution statement

TJ Steeman: Conceptualization, Investigation, Validation, Data curation, Formal analysis, Visualization, Methodology, Writing – original draft, Writing – review & editing. AMJ Weiner: Conceptualization, Investigation, Methodology, Data curation, Writing – review & editing, Resources, Funding acquisition. JA Rubiolo: Data curation, Formal analysis, Investigation, Writing – review & editing. LE Sánchez: Supervision, Resources, Writing – review & editing. NB Calcaterra: Conceptualization, Supervision, Writing – original draft, Writing – review & editing, Funding acquisition.

## 7 Data availability

Data will be available upon request.

## 8 Acknowledgements

We thank Sebastián Graziati for expert fish care, Silvana Sut for laboratory assistance and Rodrigo Vena for microscopy assistance. TJS is a fellow of CONICET, JAR is a Staff member of CONICET and AMJW and NBC have been Staff members of CONICET and Universidad Nacional de Rosario. AMJW is currently employed at BBD BioPhenix, Spain.

## 10 Supplementary data

**Supplementary Figure S 1.**
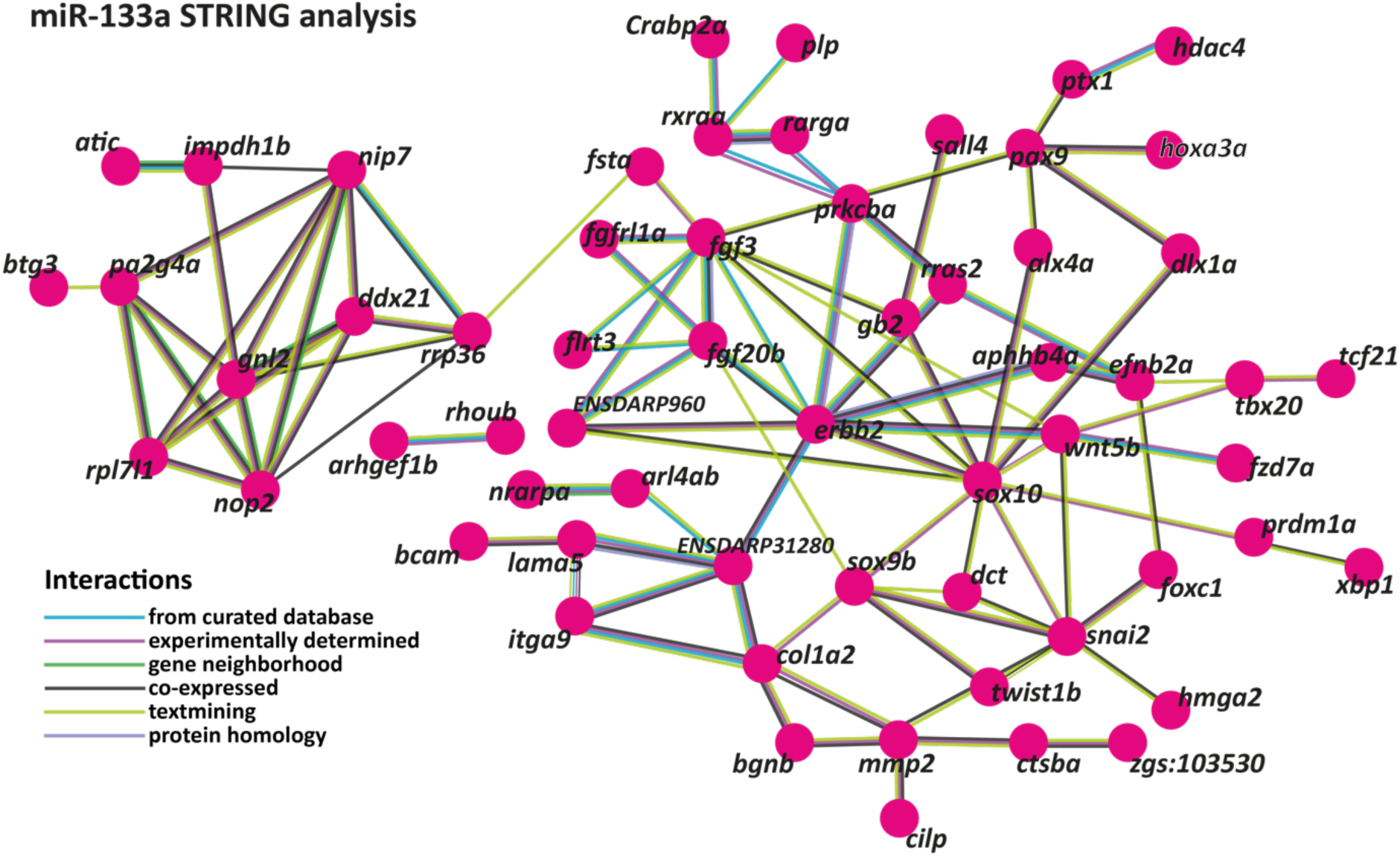
STRING interaction network for miR-133a target genes in zebrafish. The network depicts predicted and experimentally validated protein-protein interactions among genes containing putative miR-133a binding sites, as identified through bioinformatic analysis. Each node represents a protein encoded by the corresponding gene, while the connecting lines indicate interactions supported by multiple data sources.

**Supplementary Figure S 2.**
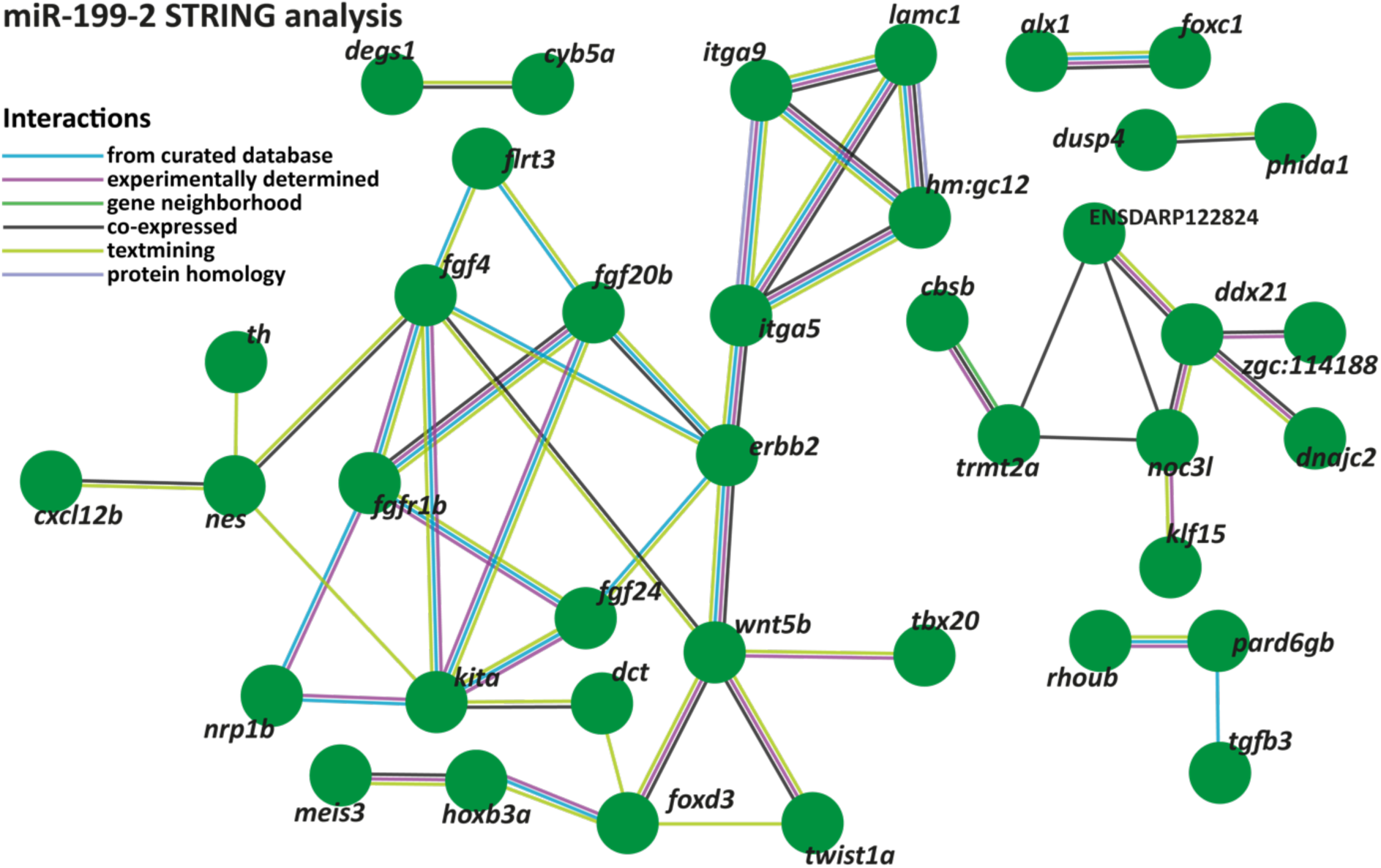
STRING interaction network for miR-1SS-2 target genes in zebrafish. The network depicts predicted and experimentally validated protein-protein interactions among genes containing putative miR-1SS-2 binding sites, as identified through bioinformatic analysis. Each node represents a protein encoded by the corresponding gene, while the connecting lines indicate interactions supported by multiple data sources.

**Supplementary Figure S 3.**
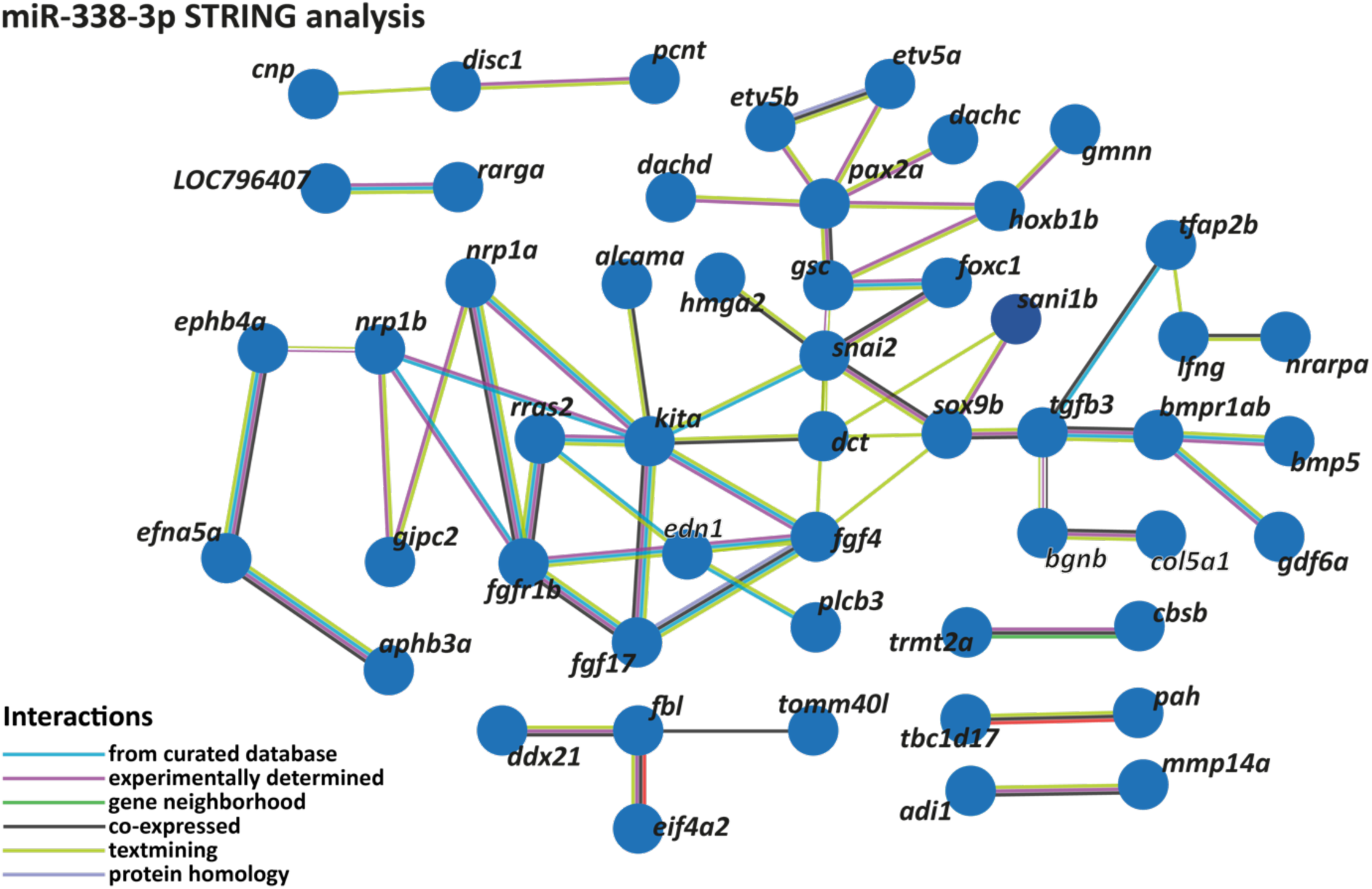
STRING interaction network for miR-338-3p target genes in zebrafish. The network depicts predicted and experimentally validated protein-protein interactions among genes containing putative miR-338-3p binding sites, as identified through bioinformatic analysis. Each node represents a protein encoded by the corresponding gene, while the connecting lines indicate interactions supported by multiple data sources.

**Supplementary Figure S 4.**
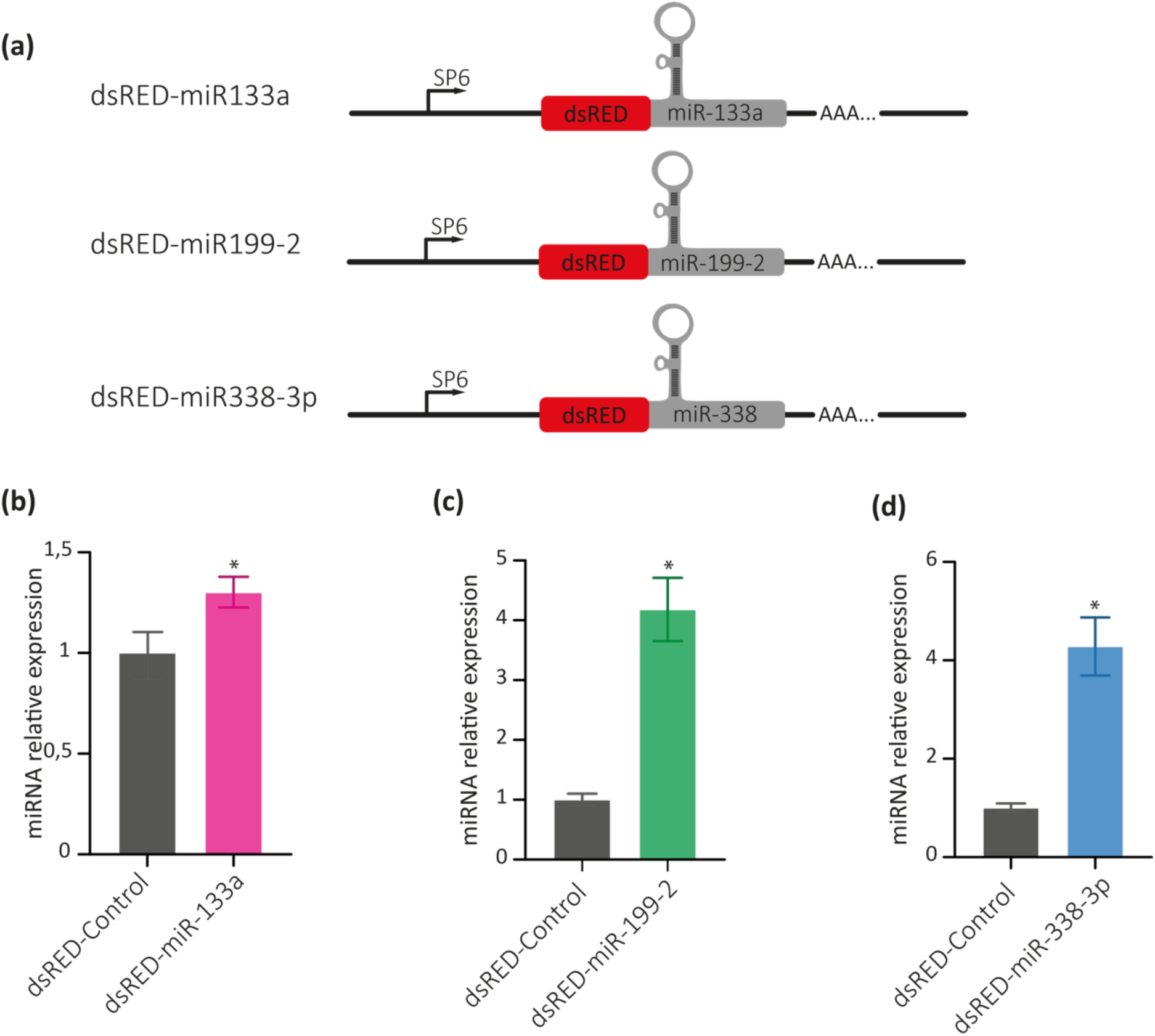
miRNA overexpression strategy. (a) Schematic representation of the constructions used to generate the synthetic dsRED/miRNA mRNAs for overexpression experiments. The red region corresponds to the dsRED coding sequence, while the grey region represents the miRNA precursor sequence. These transcripts were synthesised using SP6 RNA polymerase (binding region indicated by arrows), and a poly-A tail was included in the construct to enhance mRNA stability and facilitate dsRED translation in zebrafish cells. (b-d) RT-qPCR quantification of miR-133a (b), miR-1SS-2 (c) and miR-338-3p (d) in RNA samples from 24 hpf zebrafish embryos microinjected with either dsRED-control or miRNA overexpression transcripts. Statistical analysis was performed using a two-tailed Student’s t-test, *p ≤ 0.05.

**Supplementary Figure S 5.**
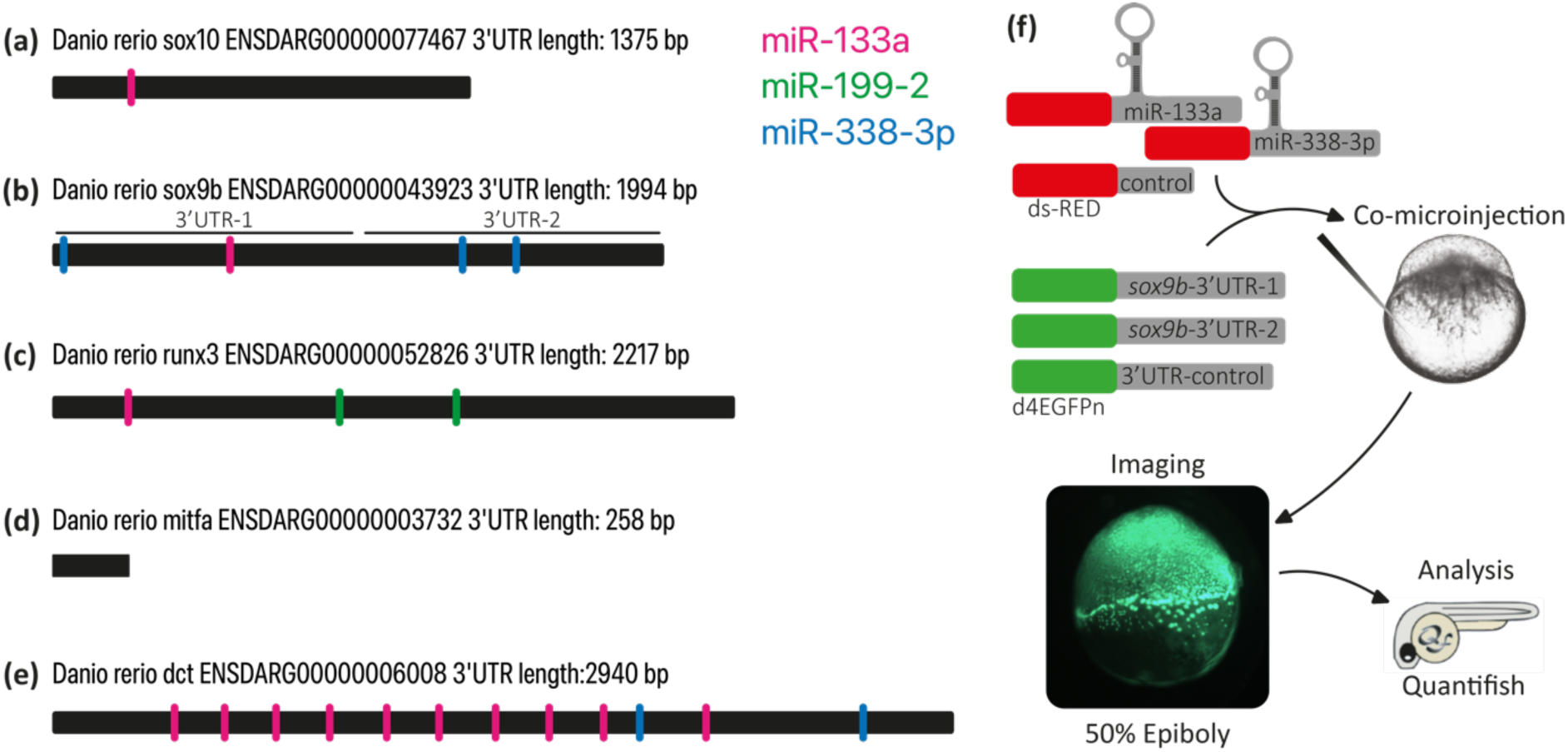
miRNA putative binding sites. (a-e) Schematic representation of the Danio rerio sox10 (a), sox9b (b), runx3 (c), mitfa (d) and dct (e) 3’UTR regions, highlighting the predicted binding sites for miR-133a (pink), miR-1SS-2 (green) and miR-338-3p (blue). The cloned fragments used in sox9b reporter assays are specifically highlighted in (b). (f) Schematic overview of the reporter assay strategy.

Supplementary Table 1: Primer sequences for cloning and RT-qPCR.

Supplementary Table 2: miRNA RNA-seq, normalized counts for GFP+ 16hpf, GFP-16hpf, GFP+ 28hpf and GFP-28hpf.

Supplementary Table S3: List of zebrafish miR-133a putative regulated genes extracted from TargetScan.org and used for Gene Ontology analyses, with context+ score.

Supplementary Table S4: Gene Ontology results for miR-133a.

Supplementary Table S5: List of zebrafish miR-199-2 putative regulated genes extracted from TargetScan.org and used for Gene Ontology analyses, with context+ score.

Supplementary Table S6: Gene Ontology results for miR-199-2.

Supplementary Table S7: List of zebrafish miR-338-3p putative regulated genes extracted from TargetScan.org and used for Gene Ontology analyses, with context+ score.

Supplementary Table S8: Gene Ontology results for miR-338-3p.

Supplementary Table S9: Zebrafish NC-related genes from early segmentation stages.

Supplementary Table S10: miRNA NC gene lists intersection.

## Bibliography

[1] Theveneau E, Mayor R. Neural crest delamination and migration: From epithelium-to-mesenchyme transition to collective cell migration. Dev Biol 2012;366:34–54. 10.1016/j.ydbio.2011.12.041.

[2] Thiery JP, Acloque H, Huang RYJ, Nieto MA. Epithelial-Mesenchymal Transitions in Development and Disease. Cell 2009;139:871–90. 10.1016/j.cell.2009.11.007.

[3] Dupin E, Le Douarin NM. The neural crest, A multifaceted structure of the vertebrates. Birth Defects Res Part C Embryo Today Rev 2014;102:187–209. 10.1002/bdrc.21080.

[4] Weiner AMJ, Coux G, Armas P, Calcaterra NB. Insights into vertebrate head development: from cranial neural crest to the modelling of neurocristopathies. Int J Dev Biol 2020. 10.1387/ijdb.200229nc.

[5] Vega-Lopez GA, Cerrizuela S, Tribulo C, Aybar MJ. Neurocristopathies: New insights 150 years after the neural crest discovery. Dev Biol 2018;444:S110–43. 10.1016/j.ydbio.2018.05.013.

[6] Martik ML, Bronner ME. Regulatory Logic Underlying Diversification of the Neural Crest. Trends Genet 2017;33:715–27. 10.1016/j.tig.2017.07.015.

[7] Hammerschmidt M, Serbedzija GN, McMahon AP. Genetic analysis of dorsoventral pattern formation in the zebrafish: requirement of a BMP-like ventralizing activity and its dorsal repressor. Genes Dev 1996;10:2452–61. 10.1101/GAD.10.19.2452.

[8] Schumacher JA, Hashiguchi M, Nguyen VH, Mullins MC. An intermediate level of BMP signaling directly specifies cranial neural crest progenitor cells in zebrafish. PloS One 2011;6. 10.1371/JOURNAL.PONE.0027403.

[9] Lewis JL, Bonner J, Modrell M, Ragland JW, Moon RT, Dorsky RI, et al. Reiterated Wnt signaling during zebrafish neural crest development. Dev Camb Engl 2004;131:1299–308. 10.1242/DEV.01007.

[10] Ferronha T, Angeles Rabadán M, Gil-Guiñon E, Le Dréau G, de Torres C, Martí E. LMO4 is an essential cofactor in the Snail2-mediated epithelial-to-mesenchymal transition of neuroblastoma and neural crest cells. J Neurosci 2013;33:2773–83. 10.1523/JNEUROSCI.4511-12.2013.

[11] Cano A, Pérez-Moreno MA, Rodrigo I, Locascio A, Blanco MJ, Del Barrio MG, et al. The transcription factor Snail controls epithelial-mesenchymal transitions by repressing E-cadherin expression. Nat Cell Biol 2000;2:76–83. 10.1038/35000025.

[12] Kelsh RN, Brand M, Jiang Y-J, Heisenberg C-P, Lin S, Haffler P, et al. Zebrafish pigmentation mutations and the processes of neural crest development. Development 1996;123:369–89. 10.1242/dev.123.1.369.

[13] Bharti K, Nguyen MTT, Skuntz S, Bertuzzi S, Arnheiter H. The other pigment cell: specification and development of the pigmented epithelium of the vertebrate eye. Pigment Cell Res 2006;19:380–94. 10.1111/J.1600-0749.2006.00318.X.

[14] Greenhill ER, Rocco A, Vibert L, Nikaido M, Kelsh RN. An Iterative Genetic and Dynamical Modelling Approach Identifies Novel Features of the Gene Regulatory Network Underlying Melanocyte Development. PLoS Genet 2011;7:e1002265. 10.1371/journal.pgen.1002265.

[15] Gur D, Bain EJ, Johnson KR, Aman AJ, Pasoili HA, Flynn JD, et al. In situ differentiation of iridophore crystallotypes underlies zebrafish stripe patterning. Nat Commun 2020 111 2020;11:1–14. 10.1038/s41467-020-20088-1.

[16] Petratou K, Subkhankulova T, Lister JA, Rocco A, Schwetlick H, Kelsh RN. A systems biology approach uncovers the core gene regulatory network governing iridophore fate choice from the neural crest. PLOS Genet 2018;14:e1007402. 10.1371/journal.pgen.1007402.

[17] Kaller M, Hermeking H. Interplay between transcription factors and microRNAs regulating epithelial-mesenchymal transitions in colorectal cancer. Adv Exp Med Biol 2016;937:71–92. 10.1007/978-3-319-42059-2_4.

[18] Lopes SS, Yang X, Müller J, Carney TJ, McAdow AR, Rauch GJ, et al. Leukocyte tyrosine kinase functions in pigment cell development. PLoS Genet 2008;4. 10.1371/JOURNAL.PGEN.1000026.

[19] Kimmel CB, Miller CT, Moens CB. Specification and morphogenesis of the zebrafish larval head skeleton. Dev Biol 2001;233:239–57. 10.1006/dbio.2001.0201.

[20] Glasauer SMK, Neuhauss SCF. Whole-genome duplication in teleost fishes and its evolutionary consequences. Mol Genet Genomics 2014;289:1045–60. 10.1007/s00438-014-0889-2.

[21] Yan Y, Willoughby J, Liu D, Crump JG, Wilson C, Miller CT, et al. A pair of Sox: distinct and overlapping functions of zebrafish sox9 co-orthologs in craniofacial and pectoral fin development. Dev Camb Engl 2005;132:1069–83. 10.1242/dev.01674.

[22] Chiang EF-L, Pai C-I, Wyatt M, Yan Y-L, Postlethwait J, Chung B. Two Sox9 Genes on Duplicated Zebrafish Chromosomes: Expression of Similar Transcription Activators in Distinct Sites. Dev Biol 2001;231:149–63. 10.1006/dbio.2000.0129.

[23] Dalcq J, Pasque V, Ghaye A, Larbuisson A, Motte P, Martial JA, et al. RUNX3, EGR1 and SOX9B Form a Regulatory Cascade Required to Modulate BMP-Signaling during Cranial Cartilage Development in Zebrafish. PLoS ONE 2012;7:e50140. 10.1371/journal.pone.0050140.

[24] Weiner AMJ. MicroRNAs and the neural crest: From induction to differentiation. Mech Dev 2018;154:98–106. 10.1016/j.mod.2018.05.009.

[25] Wienholds E, Koudijs MJ, van Eeden FJM, Cuppen E, Plasterk RHA. The microRNA-producing enzyme Dicer1 is essential for zebrafish development. Nat Genet 2003;35:217–8. 10.1038/ng1251.

[26] Kloosterman WP, Steiner FA, Berezikov E, de Bruijn E, van de Belt J, Verheul M, et al. Cloning and expression of new microRNAs from zebrafish. Nucleic Acids Res 2006;34:2558–69. 10.1093/nar/gkl278.

[27] Bartel DP. MicroRNAs: Target Recognition and Regulatory Functions. Cell 2009;136:215–33. 10.1016/j.cell.2009.01.002.

[28] Lewis BP, Burge CB, Bartel DP. Conserved Seed Pairing, Often Flanked by Adenosines, Indicates that Thousands of Human Genes are MicroRNA Targets. Cell 2005;120:15–20. 10.1016/j.cell.2004.12.035.

[29] Stark A, Brennecke J, Bushati N, Russell RB, Cohen SM. Animal microRNAs confer robustness to gene expression and have a significant impact on 3ʹUTR evolution. Cell 2005;123:1133–46. 10.1016/j.cell.2005.11.023.

[30] Friedman RC, Farh KKH, Burge CB, Bartel DP. Most mammalian mRNAs are conserved targets of microRNAs. Genome Res 2009;19:92–105. 10.1101/GR.082701.108.

[31] Giraldez AJ, Cinalli RM, Glasner ME, Enright AJ, Thomson JM, Baskerville S, et al. MicroRNAs regulate brain morphogenesis in zebrafish. Science 2005;308:833–8. 10.1126/science.1109020.

[32] Giraldez AJ, Mishima Y, Rihel J, Grocock RJ, Dongen SV, Inoue K, et al. Zebrafish MiR-430 Promotes Deadenylation and Clearance of Maternal mRNAs. Sci New Ser 2006;312:75–9.

[33] Antonaci M, Wheeler GN. MicroRNAs in neural crest development and neurocristopathies. Biochem Soc Trans 2022;50:965–74. 10.1042/BST20210828.

[34] Bamforth SD, Burn J. DiGeorge Syndrome. Brenners Encycl Genet Second Ed 2023:319–21. 10.1016/B978-0-12-374984-0.00402-2.

[35] Syeda ZA, Langden SSS, Munkhzul C, Lee M, Song SJ. Regulatory Mechanism of MicroRNA Expression in Cancer. Int J Mol Sci 2020 Vol 21 Page 1723 2020;21:1723. 10.3390/IJMS21051723.

[36] Svoronos AA, Engelman DM, Slack FJ. OncomiR or tumor suppressor? The duplicity of MicroRNAs in cancer. Cancer Res 2016;76:3666–70. 10.1158/0008-5472.CAN-16-0359/660572/P/ONCOMIR-OR-TUMOR-SUPPRESSOR-THE-DUPLICITY-OF.

[37] Kwak J, Park OK, Jung YJ, Hwang BJ, Kwon SH, Kee Y. Live image profiling of neural crest lineages in zebrafish transgenic lines. Mol Cells 2013;35:255–60. 10.1007/s10059-013-0001-5.

[38] Avellino R, Carrella S, Pirozzi M, Risolino M, Salierno FG, Franco P, et al. miR-204 targeting of Ankrd13A controls both mesenchymal neural crest and lens cell migration. PloS One 2013;8. 10.1371/JOURNAL.PONE.0061099.

[39] Gessert S, Bugner V, Tecza A, Pinker M, Kühl M. FMR1/FXR1 and the miRNA pathway are required for eye and neural crest development. Dev Biol 2010;341:222–35. 10.1016/J.YDBIO.2010.02.031.

[40] Chen PY, Manninga H, Slanchev K, Chien M, Russo JJ, Ju J, et al. The developmental miRNA profiles of zebrafish as determined by small RNA cloning. Genes Dev 2005;19:1288–93. 10.1101/gad.1310605.

[41] Wienholds E, Kloosterman WP, Miska E, Alvarez-Saavedra E, Berezikov E, de Bruijn E, et al. MicroRNA Expression in Zebrafish Embryonic Development. Science 2005;309:310–1. 10.1126/science.1114519.

[42] Lagos-Quintana M, Rauhut R, Yalcin A, Meyer J, Lendeckel W, Tuschl T. Identification of tissue-specific MicroRNAs from mouse. Curr Biol 2002;12:735–9. 10.1016/S0960-9822(02)00809-6.

[43] Boštjančič E, Zidar N, Štajer D, Glavač D. MicroRNAs miR-1, miR-133a, miR-133b and miR-208 are dysregulated in human myocardial infarction. Cardiology 2010;115:163–9. 10.1159/000268088.

[44] Castoldi G, di Gioia CRT, Bombardi C, Catalucci D, Corradi B, Gualazzi MG, et al. MiR-133a regulates collagen 1A1: Potential role of miR-133a in myocardial fibrosis in angiotensin II-dependent hypertension. J Cell Physiol 2012;227:850–6. 10.1002/jcp.22939.

[45] Yin VP, Lepilina A, Smith A, Poss KD. Regulation of zebrafish heart regeneration by miR-133. Dev Biol 2012;365:319–27. 10.1016/j.ydbio.2012.02.018.

[46] Koutsoulidou A, Mastroyiannopoulos NP, Furling D, Uney JB, Phylactou LA. Expression of miR-1, miR-133a, miR-133b and miR-206 increases during development of human skeletal muscle. BMC Dev Biol 2011;11:1–9. 10.1186/1471-213X-11-34.

[47] Wu Z sheng, Wang C qun, Xiang R, Liu X, Ye S, Yang X qing, et al. Loss of miR-133a expression associated with poor survival of breast cancer and restoration of miR-133a expression inhibited breast cancer cell growth and invasion. BMC Cancer 2012;12:1–10. 10.1186/1471-2407-12-51.

[48] He B, Xiao J, Ren AJ, Zhang YF, Zhang H, Chen M, et al. Role of miR-1 and miR-133a in myocardial ischemic postconditioning. J Biomed Sci 2011;18:1–10. 10.1186/1423-0127-18-22.

[49] Kawakami K, Enokida H, Chiyomaru T, Tatarano S, Yoshino H, Kagara I, et al. The functional significance of miR-1 and miR-133a in renal cell carcinoma. Eur J Cancer 2012;48:827–36. 10.1016/j.ejca.2011.06.030.

[50] Kojima S, Chiyomaru T, Kawakami K, Yoshino H, Enokida H, Nohata N, et al. Tumour suppressors miR-1 and miR-133a target the oncogenic function of purine nucleoside phosphorylase (PNP) in prostate cancer. Br J Cancer 2012;106:405–13. 10.1038/bjc.2011.462.

[51] Rao PK, Missiaglia E, Shields L, Hyde G, Yuan B, Shepherd CJ, et al. Distinct roles for miR-1 and miR-133a in the proliferation and differentiation of rhabdomyosarcoma cells. FASEB J 2010;24:3427–37. 10.1096/fj.09-150698.

[52] Yoshino H, Chiyomaru T, Enokida H, Kawakami K, Tatarano S, Nishiyama K, et al. The tumour-suppressive function of miR-1 and miR-133a targeting TAGLN2 in bladder cancer. Br J Cancer 2011;104:808–18. 10.1038/bjc.2011.23.

[53] Li M, Ding W, Tariq MA, Chang W, Zhang X, Xu W, et al. A circular transcript of ncx1 gene mediates ischemic myocardial injury by targeting miR-133a-3p. Theranostics 2018;8:5855–69. 10.7150/thno.27285.

[54] Li H, Yang J, Wei X, Song C, Dong D, Huang Y, et al. CircFUT10 reduces proliferation and facilitates differentiation of myoblasts by sponging miR-133a. J Cell Physiol 2018;233:4643–51. 10.1002/jcp.26230.

[55] Lim LP, Glasner ME, Yekta S, Burge CB, Bartel DP. Vertebrate microRNA genes. Science 2003;299:1540. 10.1126/science.1080372.

[56] Koshizuka K, Hanazawa T, Kikkawa N, Arai T, Okato A, Kurozumi A, et al. Regulation of ITGA3 by the anti-tumor miR-199 family inhibits cancer cell migration and invasion in head and neck cancer. Cancer Sci 2017;108:1681–92. 10.1111/cas.13298.

[57] Zhang HY, Li CH, Wang XC, Luo YQ, Cao XD, Chen JJ. MiR-199 inhibits EMT and invasion of hepatoma cells through inhibition of Snail expression. Eur Rev Med Pharmacol Sci 2019;23:7884–91. 10.26355/eurrev_201909_18998.

[58] Wang W, Guo Z, Yang S, Wang H, Ding W. Upregulation of miR-199 attenuates TNF-α-induced Human nucleus pulposus cell apoptosis by downregulating MAP3K5. Biochem Biophys Res Commun 2018;505:917–24. 10.1016/j.bbrc.2018.09.194.

[59] Kim J, Krichevsky A, Grad Y, Hayes GD, Kosik KS, Church GM, et al. Identification of many microRNAs that copurify with polyribosomes in mammalian neurons. Proc Natl Acad Sci 2004;101:360–5. 10.1073/pnas.2333854100.

[60] Howe JR, Li ES, Streeter SE, Rahme GJ, Chipumuro E, Russo GB, et al. MIR-338-3p regulates neuronal maturation and suppresses glioblastoma proliferation. PLoS ONE 2017;12:1–21. 10.1371/journal.pone.0177661.

[61] Chen X, Pan M, Han L, Lu H, Hao X, Dong Q. miR-338-3p suppresses neuroblastoma proliferation, invasion and migration through targeting PREX2a. FEBS Lett 2013;587:3729–37. 10.1016/j.febslet.2013.09.044.

[62] Huang N, Wu Z, Lin L, Zhou M, Wang L, Ma H, et al. MiR-338-3p inhibits epithelial-mesenchymal transition in gastric cancer cells by targeting ZEB2 and MACC1/Met/Akt signaling. Oncotarget 2015;6:15222–34. 10.18632/oncotarget.3835.

[63] Liu C, Wang Z, Wang Y, Gu W. miR-338 suppresses the growth and metastasis of OSCC cells by targeting NRP1. Mol Cell Biochem 2015;398:115–22. 10.1007/s11010-014-2211-3.

[64] Huang XH, Chen JS, Wang Q, Chen XL, Wen L, Chen LZ, et al. MiR-338-3p suppresses invasion of liver cancer cell by targeting smoothened. J Pathol 2011;225:463–72. 10.1002/path.2877.

[65] Moreno P, Fexova S, George N, Manning JR, Miao Z, Mohammed S, et al. Expression Atlas update: gene and protein expression in multiple species. Nucleic Acids Res 2022;50:D129–40. 10.1093/nar/gkab1030.

[66] White RJ, Collins JE, Sealy IM, Wali N, Dooley CM, Digby Z, et al. A high-resolution mRNA expression time course of embryonic development in zebrafish. eLife 2017;6. 10.7554/eLife.30860.

[67] Vastenhouw NL, Cao WX, Lipshitz HD. The maternal-to-zygotic transition revisited. Development 2019;146:dev161471. 10.1242/dev.161471.

[68] Bell D, Leung K, Wheatley S, Ng L, Zhou S, Ling K, et al. SOX9 directly regulates the type-II collagen gene. Nat Genet 1997;16:174–8.

[69] Lefebvre V, Li P, De Crombrugghe B. A new long form of Sox5 (L-Sox5), Sox6 and Sox9 are coexpressed in chondrogenesis and cooperatively activate the type II collagen gene. EMBO J 1998;17:5718–33. 10.1093/EMBOJ/17.19.5718.

[70] Townley AK, Feng Y, Schmidt K, Carter DA, Porter R, Verkade P, et al. Efficient coupling of Sec23-Sec24 to Sec13-Sec31 drives COPII-dependent collagen secretion and is essential for normal craniofacial development. J Cell Sci 2008;121:3025–34. 10.1242/JCS.031070.

[71] Xu P, Balczerski B, Ciozda A, Louie K, Oralova V, Huysseune A, et al. Fox proteins are modular competency factors for facial cartilage and tooth specification. Dev Camb 2018;145. 10.1242/DEV.165498/264659/AM/FOX-PROTEINS-ARE-MODULAR-COMPETENCY-FACTORS-FOR.

[72] Kelsh RN, Raible DW. Specification of Zebrafish Neural Crest. Results Probl. Cell Differ., vol. 40, 2002, p. 216–36. 10.1007/978-3-540-46041-1_11.

[73] Guglielmi L, Bühler A, Moro E, Argenton F, Poggi L, Carl M. Temporal control of Wnt signaling is required for habenular neuron diversity and brain asymmetry. Dev Camb 2020;147. 10.1242/dev.182865.

[74] Yin A, Korzh V, Gong Z. Perturbation of zebrafish swimbladder development by enhancing Wnt signaling in Wif1 morphants. Biochim Biophys Acta - Mol Cell Res 2012;1823:236–44. 10.1016/j.bbamcr.2011.09.018.

[75] Santoriello C, Sporrij A, Yang S, Flynn RA, Henriques T, Dorjsuren B, et al. RNA helicase DDX21 mediates nucleotide stress responses in neural crest and melanoma cells. Nat Cell Biol 2020;22:372–9. 10.1038/s41556-020-0493-0.

[76] Johansson JA, Marie KL, Lu Y, Brombin A, Santoriello C, Zeng Z, et al. PRL3-DDX21 Transcriptional Control of Endolysosomal Genes Restricts Melanocyte Stem Cell Differentiation. Dev Cell 2020;54:317–332.e9. 10.1016/j.devcel.2020.06.013.

[77] Gallardo VE, Behra M. Fluorescent activated cell sorting (FACS) combined with gene expression microarrays for transcription enrichment profiling of zebrafish lateral line cells. Methods 2013;62:226–31. 10.1016/j.ymeth.2013.06.005.

[78] Kloosterman WP, Plasterk RHA. The Diverse Functions of MicroRNAs in Animal Development and Disease. Dev Cell 2006;11:441–50. 10.1016/j.devcel.2006.09.009.

[79] Kim VN. MicroRNA biogenesis: coordinated cropping and dicing. Nat Rev Mol Cell Biol 2005;6:376–85. 10.1038/nrm1644.

[80] Gilbert SF. Developmental Biology. 2000.

[81] Giovannini C, Fornari F, Dallo R, Gagliardi M, Nipoti E, Vasuri F, et al. MiR-199-3p replacement affects E-cadherin expression through Notch1 targeting in hepatocellular carcinoma. Acta Histochem 2018;120:95–102. 10.1016/j.acthis.2017.12.004.

[82] Schmidt R, Strähle U, Scholpp S. Neurogenesis in zebrafish – from embryo to adult. Neural Develop 2013;8:3. 10.1186/1749-8104-8-3.

[83] Flores MV, Lam EYN, Crosier P, Crosier K. A hierarchy of Runx transcription factors modulate the onset of chondrogenesis in craniofacial endochondral bones in zebrafish. Dev Dyn 2006;235:3166–76. 10.1002/dvdy.20957.

[84] Nordlund JJ, Boissy RE, Hearing VJ, King RA, Oetting WS, Ortonne JP. The Pigmentary System: Physiology and Pathophysiology: Second Edition. Pigment Syst Physiol Pathophysiol Second Ed 2007:1–1229. 10.1002/9780470987100.

[85] Sánchez-Vásquez E, Bronner ME, Strobl-Mazzulla PH. Epigenetic inactivation of miR-203 as a key step in neural crest epithelial-to-mesenchymal transition. Development 2019;146:dev171017. 10.1242/dev.171017.

[86] Yu X, Odenthal M, Fries JWU. Exosomes as miRNA Carriers: Formation–Function– Future. Int J Mol Sci 2016 Vol 17 Page 2028 2016;17:2028. 10.3390/IJMS17122028.

[87] Lukiw WJ, Pogue AI. Vesicular Transport of Encapsulated microRNA between Glial and Neuronal Cells. Int J Mol Sci 2020 Vol 21 Page 5078 2020;21:5078. 10.3390/IJMS21145078.

[88] Xu L, Yang BF, Ai J. MicroRNA transport: A new way in cell communication. J Cell Physiol 2013;228:1713–9. 10.1002/JCP.24344.

[89] Westerfield M. A guide for the laboratory use of zebrafish (Danio rerio). Zebrafish Book 2000;4.

[90] Kimmel CB, Ballard WW, Kimmel SR, Ullmann B, Schilling TF. Stages of embryonic development of the zebrafish. Dev Dyn 1995;203:253–310. 10.1002/aja.1002030302.

[91] Langmead B, Trapnell C, Pop M, Salzberg SL. Ultrafast and memory-efficient alignment of short DNA sequences to the human genome. Genome Biol 2009;10:R25. 10.1186/gb-2009-10-3-r25.

[92] Putri GH, Anders S, Pyl PT, Pimanda JE, Zanini F. Analysing high-throughput sequencing data in Python with HTSeq 2.0. Bioinformatics 2022;38:2943–5. 10.1093/bioinformatics/btac166.

[93] Love MI, Huber W, Anders S. Moderated estimation of fold change and dispersion for RNA-seq data with DESeq2. Genome Biol 2014;15:550. 10.1186/s13059-014-0550-8.

[94] Ulitsky I, Shkumatava A, Jan CH, Subtelny AO, Koppstein D, Bell GW, et al. Extensive alternative polyadenylation during zebrafish development. Genome Res 2012;22:2054–66. 10.1101/gr.139733.112.

[95] Huang DW, Sherman BT, Lempicki RA. Systematic and integrative analysis of large gene lists using DAVID bioinformatics resources. Nat Protoc 2009;4:44–57. 10.1038/nprot.2008.211.

[96] Mi H, Muruganujan A, Ebert D, Huang X, Thomas PD. PANTHER version 14: more genomes, a new PANTHER GO-slim and improvements in enrichment analysis tools. Nucleic Acids Res 2019;47:D419–26. 10.1093/nar/gky1038.

[97] Raudvere U, Kolberg L, Kuzmin I, Arak T, Adler P, Peterson H, et al. g:Profiler: a web server for functional enrichment analysis and conversions of gene lists (2019 update). Nucleic Acids Res 2019;47:W191–8. 10.1093/nar/gkz369.

[98] Kramer MF. Stem-Loop RT-qPCR for miRNAs. Curr Protoc Mol Biol 2011;95:15.10.1-15.10.15. 10.1002/0471142727.mb1510s95.

[99] Pfaffl MW, Horgan GW, Dempfle L. Relative expression software tool (REST©) for group-wise comparison and statistical analysis of relative expression results in real-time PCR. vol. 30. 2002.

[100] Bustin SA, Benes V, Garson JA, Hellemans J, Huggett J, Kubista M, et al. The MIQE Guidelines: Minimum Information for Publication of Quantitative Real-Time PCR Experiments. Clin Chem 2009;55:611–22. 10.1373/clinchem.2008.112797.

[101] Steeman TJ, Rubiolo JA, Sánchez LE, Calcaterra NB, Weiner AMJ. Conservation of Zebrafish MicroRNA-145 and Its Role during Neural Crest Cell Development. Genes 2021;12:1023. 10.3390/genes12071023.

[102] Stirling DR, Suleyman O, Gil E, Elks PM, Torraca V, Noursadeghi M, et al. Analysis tools to quantify dissemination of pathology in zebrafish larvae. Sci Rep 2020;10:3149. 10.1038/s41598-020-59932-1.

[103] Weiner AMJ, Scampoli NL, Steeman TJ, Dooley CM, Busch-Nentwich EM, Kelsh RN, et al. Dicer1 is required for pigment cell and craniofacial development in zebrafish. Biochim Biophys Acta BBA - Gene Regul Mech 2019;1862:472–85. 10.1016/j.bbagrm.2019.02.005.

[104] Schneider CA, Rasband WS, Eliceiri KW. NIH Image to ImageJ: 25 years of image analysis. Nat Methods 2012;9:671–5. 10.1038/nmeth.2089.

